# Experience-dependent changes in hippocampal spatial activity and hippocampal circuit function are disrupted in a rat model of Fragile X Syndrome

**DOI:** 10.1101/2021.09.17.460768

**Authors:** Antonis Asiminas, Sam A Booker, Owen R Dando, Zrinko Kozic, Daisy Arkell, Felicity H Inkpen, Anna Sumera, Irem Akyel, Peter C Kind, Emma R Wood

## Abstract

Fragile X syndrome (FXS) is a common single gene cause of intellectual disability and Autism Spectrum Disorder. Cognitive inflexibility is one of the hallmarks of FXS with affected individuals showing extreme difficulty adapting to novel or complex situations. To explore the neural correlates of this cognitive inflexibility, we used a rat model of FXS (*Fmr1*^-/y^), and recorded from the CA1 region of the hippocampus while animals habituated in a novel environment for two consecutive days. On the first day of exploration, the firing rate and spatial tuning of CA1 pyramidal neurons was similar between wild-type (WT) and *Fmr1*^-/y^ rats. However, while CA1 pyramidal neurons from WT rats showed experience-dependent changes in firing and spatial tuning between the first and second day of exposure to the environment, these changes were decreased or absent in CA1 neurons of *Fmr1*^-/y^ rats. These findings were consistent with increased excitability of *Fmr1*^-/y^ CA1 neurons in *ex-vivo* hippocampal slices, which correlated with reduced synaptic inputs from the medial entorhinal cortex. Lastly, activity patterns of CA1 pyramidal neurons were discoordinated with respect to hippocampal oscillatory activity in *Fmr1*^-/y^ rats. These findings suggest a network-level origin of cognitive deficits in FXS.

## Introduction

Fragile X Syndrome (FXS) is a genetic neurodevelopmental disorder caused by a CGG repeat expansion in the promoter region of the *FMR1* gene, resulting in its epigenetic silencing and subsequent loss of its protein product FMRP (Colak et al., 2014; Pieretti et al., 1991; Verkerk et al., 1991). Although FXS is a rare disorder, it is one of the most commonly inherited forms of intellectual disability (ID) and autism spectrum disorder (ASD) (Coffee et al., 2009; Hunter et al., 2014). FMRP has numerous functions from translational repression of a wide variety of mRNAs (Darnell et al., 2011) to directly binding ion channels to regulate excitability (Deng et al., 2013; Zhang et al., 2012). Hence, elucidating direct mechanistic links between FMRP loss and specific behavioural phenotypes has been challenging. Bridging this gap requires an understanding of the functional changes to neural circuits underlying behaviour resulting from the absence FMRP.

Rodent models of FXS have been instrumental in understanding how FMRP loss leads to behavioural and cognitive changes. *Fmr1*^-/y^ mice exhibit deficits in a range of cognitive tasks as well as synaptic plasticity abnormalities in various brain areas (Sidorov, Auerbach & Bear, 2013; Dahlhaus, 2018). Hippocampal circuits are affected in both FXS patients and rodent models of FXS (Bostrom et al., 2016) and hippocampal pathophysiology has been well-characterised in rodent models. Studies in *Fmr1*^-/y^ mice have identified changes in CA1 pyramidal neuron excitability (Booker et al., 2020, Ordemann et al., 2021, Luque et al., 2018) and the strength of inputs they receive (Booker et al., 2020), alterations in hippocampal synaptic plasticity (Bhakar, Dölen & Bear 2012; Bostrom et al., 2016; Huber et al., 2002) and deficits in some hippocampus-dependent learning and memory tasks (Bostrom et al., 2016; Santos, Kanellopoulos & Bagni, 2014). Consistent with these findings, we have reported deficits in a subset of hippocampus-dependent memory tasks and disturbances in hippocampal plasticity in *Fmr1*^-/y^ rat models of FXS (Asiminas et al., 2019; Till et al., 2015).

Recent studies have begun to shed light on the effects of FMRP loss on hippocampal function, using *in vivo* electrophysiological recordings from the hippocampus of freely moving mice. The basic firing properties of hippocampal pyramidal neurons of *Fmr1*^-/y^ mice remain largely normal, but analysis of the temporal coordination of spiking activity and local field potentials (LFP) point to disrupted circuit organization and function (Arbab et al., 2018; Arbab, Pennartz, & Battaglia, 2018; Boone et al., 2018; Dvorak et al., 2018; Radwan, Dvorak, & Fenton, 2016; Talbot et al., 2018). Furthermore, *ex-vivo* studies demonstrated that environmental novelty can induce exaggerated changes in hippocampal synaptic function of *Fmr1*^-/y^ mice (Talbot et al., 2018).

A well characterized feature of hippocampal pyramidal neurons is their spatially modulated firing, such that a given neuron (place cell) fires when the animal moves through a specific region of the environment (the cell’s place field; O’Keefe & Dostrovsky, 1971). The activity of place cells is crucial both for spatial navigation, and for the encoding and consolidation of new spatial and episodic memories (Eichenbaum et al., 1999). The encoding of novel environments by hippocampal place cells is a fundamental learning process that is supported by cellular mechanisms involved in long-term plasticity (Moser, Rowland & Moser, 2015). Thus, studying the firing pattern dynamics of place cells in novel environments provides a powerful system for probing hippocampal correlates of memory formation. CA1 pyramidal neurons form place fields during an animal’s first exposure to a novel environment, but the spatial tuning and stability of their fields improve as a function of repeated exposure, and these changes are dependent on plasticity processes within the hippocampal formation (Bett et al., 2013; Bostock, Muller, & Kubie, 1991; Cacucci et al., 2007; Frank, 2004; Kentros et al., 1998; Lever et al., 2002; O’Keefe & Dostrovsky, 1971; Quirk, Muller, & Kubie, 1990).

The aim of the current study was to examine hippocampal pyramidal neuron activity as rats habituate to a novel environment. Building upon our previous findings, that *Fmr1*^-/y^ rats exhibit hippocampal plasticity abnormalities and deficits in hippocampus-dependent associative recognition memory and the reduced cognitive flexibility in individuals with FXS, we predicted that experience-dependent changes in the spatial tuning and the spatial stability of hippocampal representations in novel environments would be disrupted in *Fmr1*^-/y^ rats.

We recorded from the CA1 pyramidal neuron layer of the dorsal hippocampus in *Fmr1*^-/y^ and WT littermate rats as they explored an initially novel environment over three 10 min exploration sessions on each of two consecutive days. We assessed the spatial activity of neurons, as well as temporal features of hippocampal activity, to gain insights in circuit-wide effects of FMRP loss. Finally, to determine the cellular mechanism of such alteration, we performed *ex-vivo* slice recordings from CA1 pyramidal neurons to examine their intrinsic and synaptic properties.

## Results

We recorded from the dorsal CA1 of adult *Fmr1*^-/y^ (n=7) and WT rats (n=7) (10-16 weeks) as they foraged for randomly scattered cereal rewards in an initially novel cylindrical environment for three 10 min sessions (10 min ITI) over each of two consecutive days (Figure 1A-B; Figure 1 Suppl 1; Figure 1 Suppl 2). Over the duration of the experiment we recorded 288 CA1 neurons from WT and 246 from *Fmr1*^-/y^ rats that were sufficiently isolated (based on L-ratio and Isolation D; for details see Methods), were classified as pyramidal neurons (spike width >250 μs and mean firing rate <5 Hz), and were active (mean firing rate >0.1 Hz) in the environment.

**Figure 1.**
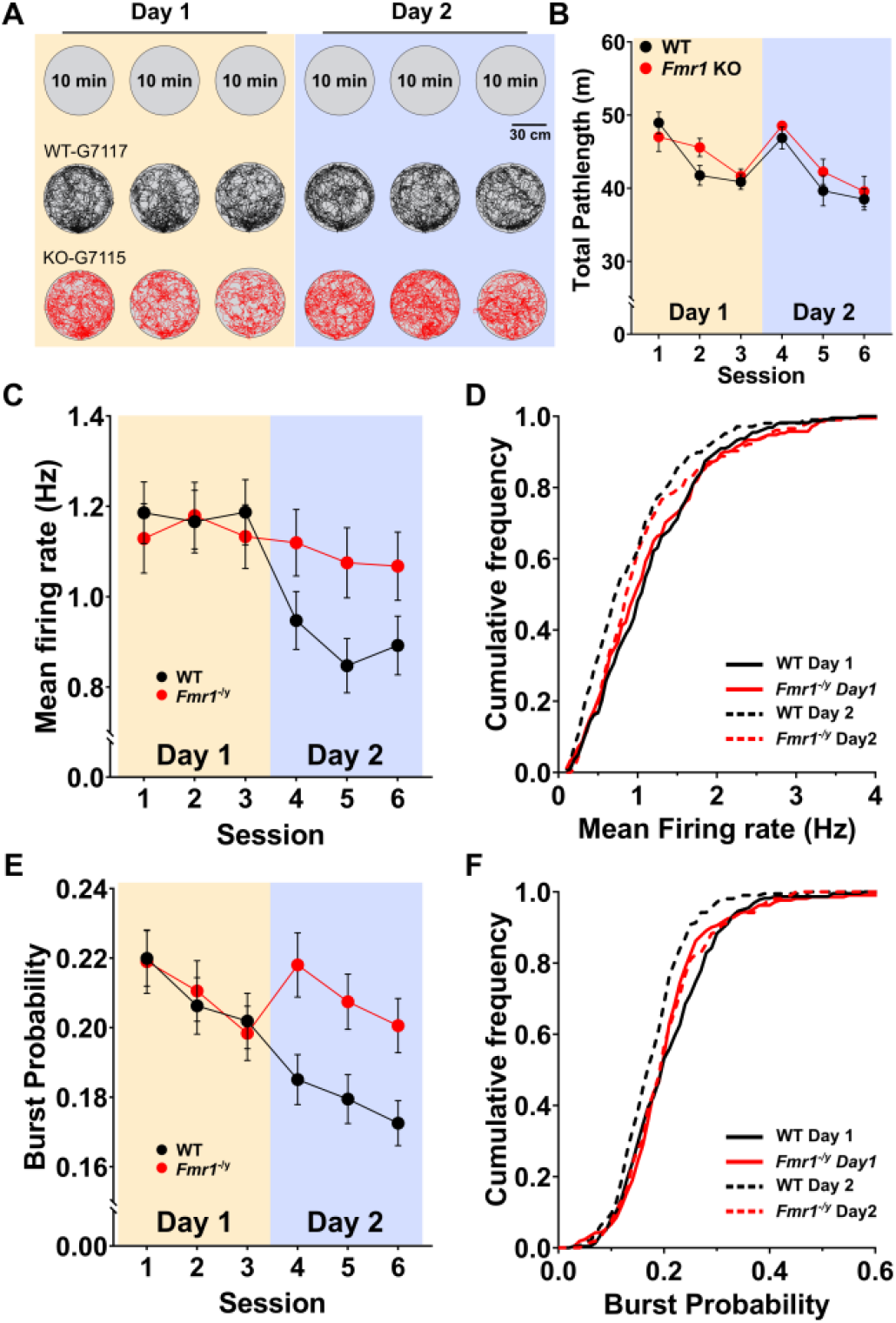
*Fmr1*^-/y^ CA1 pyramidal neurons exhibit attenuated experience-dependent decrease in mean firing rate. (**A**) Schematic of the recording protocol (Top) and example trajectories from a WT (black) and an *Fmr1*^-/y^ rat (red). (**B**) Total path length decreased across the three sessions of each day and across days for both WT and *Fmr1*^-/y^ rats. Data points depict rat means; error bars depict SEM; n_WT_ = 7, n_*Fmr1-/y*_ = 7. (**C**) Mean firing rate of CA1 pyramidal neurons decreases on the second day of exploring an initially novel environment more prominently in WT compared to *Fmr1*^*-/y*^ rats. Data represent cell means and SEMs. (**D**) Cumulative distributions of mean firing rate for unique pyramidal neurons recorded from WT and *Fmr1*^*-/y*^ rats during three sessions of Day 1 and three sessions of Day 2. For neurons identified in more than one session within a day, the mean firing rate across these sessions is included in the analysis. Both WT and *Fmr1*^*-/y*^ neuronal populations shift towards lower firing rates from Day 1 to Day 2 but WT pyramidal neurons exhibit a larger negative shift. (**E**) Burst probability of pyramidal neurons decreases over repeated exposure to a novel environment in WT, but not in *Fmr1*^*-/y*^ rats. Data represent cell means and SEMs. (**F**) Same as (**D**) for burst probability. Only WT neuronal populations shift towards lower burst probability from Day 1 to Day 2. N_WT-D1_=222, N_WT-D2_=207, N_KO-D1_=211, N_KO-D2_=205. Pale yellow and pale purple backgrounds denote data from Day 1 and Day 2 respectively.

### Experience-dependent decrease in CA1 pyramidal neuron activity is attenuated in *Fmr1*^-/y^ rats

Hippocampal pyramidal neurons typically show higher firing rates in novel than in familiar environments (Karlsson & Frank, 2008; Nitz & McNaughton, 2004) and this phenomenon has been linked to enhanced input integration (Cohen, Bolstad & Lee, 2017) during novelty. We therefore examined whether the mean firing rates of CA1 pyramidal neurons differed between WT and *Fmr1*^-/y^ rats, and whether they changed as a function of experience in the novel environment. Pyramidal neurons from WT and *Fmr1*^-/y^ rats showed very similar firing rates on Day 1 (sessions 1-3), but while the neurons from WT rats exhibited lower mean firing rates on Day 2 than Day 1, those of *Fmr1*^-/y^ rats did not differ significantly between days (Figure 1C; Figure 1 Suppl 3A&B; Figure 1 Suppl 4; Linear Mixed-Effects (LME) modelling with rats and cells as random factors: genotype x day interaction p=0.006; *posthoc* tests Day 1 vs Day 2, WT: p<0.0001; *Fmr1*^-/y^: p=0.131). However, between genotype *posthoc* tests did not reveal significant differences between WT and *Fmr1*^-/y^ on either day (Day 1: p=0.992; Day 2, p=0.130). Analysis of the distribution of firing rates for all pyramidal neurons from WT and *Fmr1*^-/y^ rats was consistent with the results from the LME modelling (Figure 1D). The distribution of firing rates for WT pyramidal neurons was significantly shifted to lower firing rates (curve shifted to the left) on Day 2 compared to Day 1, while the distribution of firing rates for *Fmr1*^-/y^ pyramidal neurons remained stable across days (Kolmogorov-Smirnov WT: D=0.1907, p<0.001; *Fmr1*^-/y^: D=0.1085, p=0.173). There was no difference in firing rate distribution between genotypes on the first day (Kolmogorov-Smirnov Day 1: D=0.0616, p=0.806), but for this analysis the distribution of mean firing rates was significantly lower in WT rats than in *Fmr1*^-/y^ rats on Day 2 (Kolmogorov-Smirnov Day 2: D=0.1481, p=0.022).

To determine the pattern of pyramidal neuron activity in WT and *Fmr1*^-/y^ rats, we calculated their burst probability, which is a well-documented property of pyramidal neurons that is controlled by perisomatic and dendritic inhibition (Royer et al., 2012). High burst probability is linked to plasticity processes underlying new place field formation (Cohen, Bolstad & Lee, 2017; Harris et al., 2001). We defined bursts in pyramidal neurons as events with more than two spikes with <10 ms inter-spike intervals (Ranck, 1973). Pyramidal neurons from WT and *Fmr1*^-/y^ rats showed similar burst probability on Day 1, and the burst probability decreased significantly between days in WT but not in *Fmr1*^-/y^ rats. (Figure 1E; Figure 1 Suppl 3C&D; Figure 1 Suppl 4; LME: genotype x day interaction p=0.005; *posthoc* tests Day 1 vs Day 2: WT: p <0.0001; *Fmr1*^-/y^ p=0.924). Similar to mean firing rates, there was no significant difference in burst probability between genotypes on either day, although there was a trend towards a difference on Day 2 (between genotype *posthoc* tests: Day 1 p=0.973, Day 2 p=0.068). There was also a significant decrease of burst probability across sessions within each day (LME: session-in-day effect p=0.013), but this did not differ between genotypes (LME: genotype x session-in-day interaction p=0.916). Analysis of the distribution of burst probabilities across all neurons revealed that only pyramidal neurons from WT rats had lower burst probabilities on Day 2 than on Day 1 (Kolmogorov-Smirnov WT: D=0.233, p<0.0001; *Fmr1*^-/y^: D=0.052, p=0.939). This analysis also indicated significantly different distributions for WT and *Fmr1*^-/y^ rats on the second day of exposure to the novel environment, but not on the first (Kolmogorov-Smirnov Day 1: D=0.123, p=0.078; Day 2: D=0.159, p=0.011) (Figure 1F).

Taken together, these analyses indicate that both the mean firing rate and burst probability of CA1 pyramidal neurons from WT and *Fmr1*^-/y^ rats is similar on the first day of exploration in a novel environment. However, on the second day of exploration, pyramidal neurons from WT rats decrease both their mean firing rate and their propensity to burst, while these experience-dependent changes in firing rate and burst probability are not seen in pyramidal neurons from *Fmr1*^-/y^ rats.

### Experience-dependent refinement of spatial tuning of CA1 pyramidal neurons is disrupted in *Fmr1*^-/y^ rats

While the mean firing rate of neurons decreases, the spatial tuning of pyramidal neurons has been shown to increase as a function of experience both in mice (Cacucci et al., 2007; Kee, Mou, Zoghbi, & Ji, 2018) and rats (Retailleau & Morris, 2018) with the biggest increase observed between the first and second day of exposure in an initially novel environment (Cacucci et al., 2007; Retailleau & Morris, 2018). We therefore assessed how the spatial firing properties of the CA1 pyramidal neurons from *Fmr1*^-/y^ and WT rats change in response to experience. The spatial tuning of a pyramidal neuron can be quantified and expressed as a number of metrics such as spatial information, sparsity and the percentage of pixels in the recording environment in which the neuron discharges. Spatial information measures the amount of information a neuron’s firing conveys about the animal’s location in an environment (Skaggs, McNaughton, Gothard, & Markus, 1993). Sparsity is a measure that reflects how compact a place field is relative to the recording arena (Jung, Wiener, McNaughton, 1994); this is closely related to the percentage of the arena in which a neuron has been active. Spatial information typically increases with repeated exposure to an environment. Consistent with previous findings, we observed an increase in the spatial information of CA1 pyramidal neurons between the first and second day of exposure to the novel environment in the WT rats. In contrast, pyramidal neurons in *Fmr1*^-/y^ rats did not show this increase (Figure 2A; Figure 2 Suppl 1A&B; Figure 2 Suppl 2; LME: genotype x day interaction p=0.005). Post-hoc comparisons of spatial information between genotypes on each day revealed no difference between genotypes on the first day in the novel environment (p=0.9304) and a non-significant trend on day two (p=0.079). There was a significant difference between the first and second day for WT neurons (p<0.001) but not for *Fmr1*^-/y^ neurons (p=0.129). Analysis of the distribution of spatial information of neurons between genotypes and days revealed a significant shift in the distribution between days for WT but not for *Fmr1*^-/y^ neurons (Kolmogorov-Smirnov WT: D=0.251, p<0.001; *Fmr1*^-/y^: D=0.081, p=0.509). Moreover, we found that the distributions of WT and *Fmr1*^-/y^ neurons did not differ on Day 1, but they did on Day 2 (Kolmogorov-Smirnov Day1: D =0.060, p=0.829; Day2: D =0.175, p=0.004)(Figure 2B).

**Figure 2.**
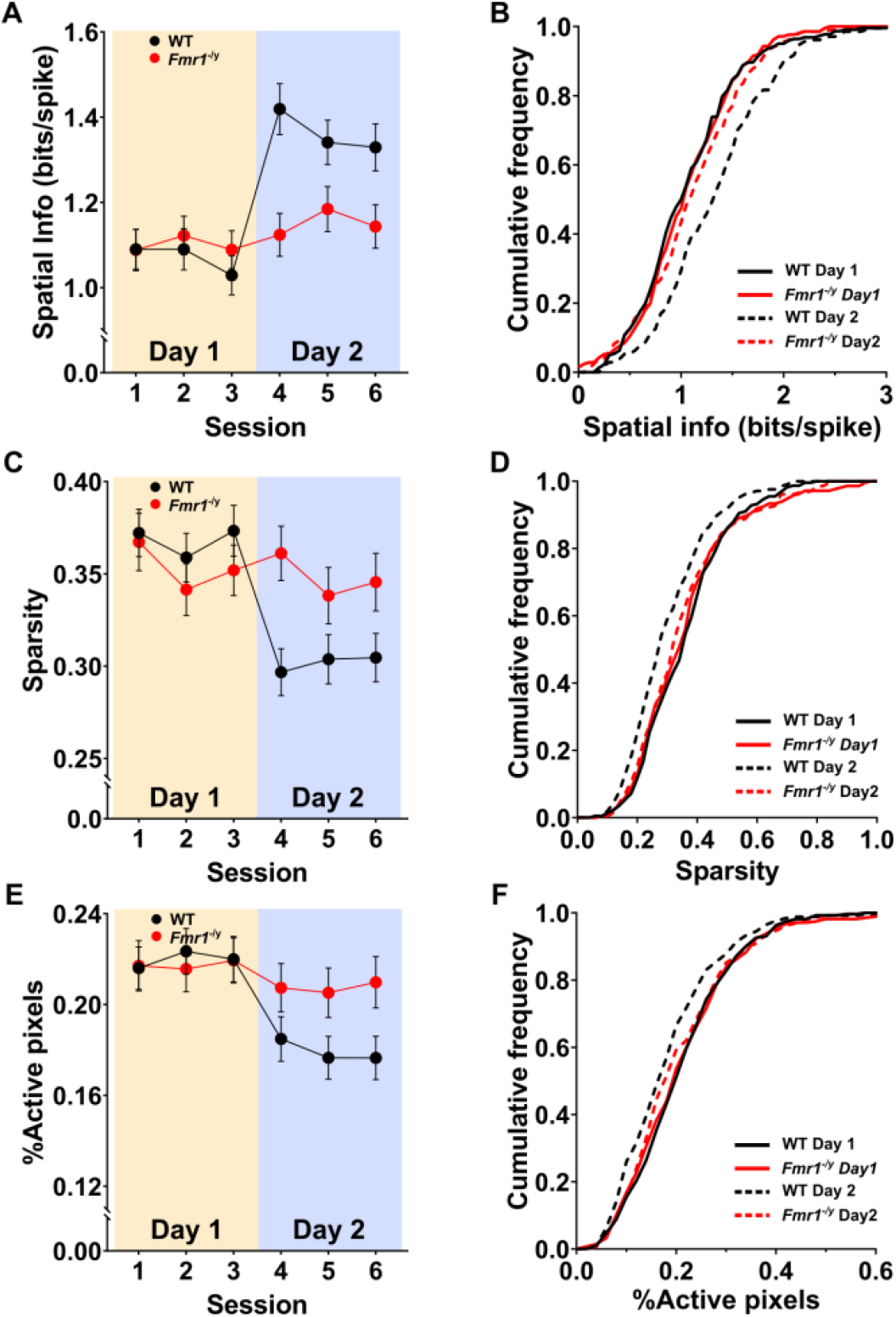
Experience-dependent refinement of spatial information coding in the CA1 pyramidal cells is impaired in *Fmr1*^*-/y*^ rats. (**A**) WT CA1 pyramidal neurons increase their spatial information over two days of exposure to the initially novel environment, while this phenomenon is not evident in *Fmr1*^*-/y*^ rats. Data represent cell means and SEMs. (**B**) Cumulative distributions of spatial information content for unique pyramidal neurons recorded from WT and *Fmr1*^*-/y*^ rats during three sessions of Day 1 and three sessions of Day 2. For neurons identified in more than one session within a day, the mean spatial information across these sessions, is included in the analysis. Only WT neuronal populations shift towards higher spatial information content from Day 1 to Day 2 (**C**) WT CA1 pyramidal neurons show trends towards a decrease of their spatial discharge sparsity over two days of exposure to an initially novel environment, while this phenomenon was not observed in *Fmr1*^*-/y*^ rats. Data represent cell means and SEMs. (**D**) Same as (**B**) for sparsity. Only WT neuronal populations shift towards lower sparsity from Day 1 to Day 2. (**E**) The proportion of the arena in which WT CA1 pyramidal neurons were active (% Active pixels) show trends towards a decrease over two days of recordings, while this phenomenon was not observed in *Fmr1*^*-/y*^ rats. Data represent cell means and SEMs. (**F**) Same as (**B**) for % Active pixels. Only WT neuronal populations shift towards lower % Active pixels from Day 1 to Day 2. N_WT-D1_=222, N_WT-D2_=207, N_KO-D1_=211, N_KO-D2_=205. Pale yellow and pale purple backgrounds denote data from Day 1 and Day 2 respectively.

The two other measures of spatial coding - sparsity and the proportion of the arena in which neurons were active (% Active pixels) - followed a very similar pattern to spatial information (Figure 2 C & E; Figure 2 Suppl 1C-F; Figure 2 Suppl 2; LME Sparsity: genotype x day interaction p=0.013; *post-hocs*, Day1: WTvsKO p=0.973, Day2: WTvsKO p=0.094, WT: Day1vsDay2 p<0.001, KO: Day1vsDay2 p=0.091; % Active pixels: genotype x day interaction p=0.070). Comparison of distributions revealed that the distributions differed between Day 1 and Day 2 for WT neurons for both measures (Kolmogorov-Smirnov Sparsity: D=0.217, p<0.001; % Active pixels: D=0.187, p=0.001) but not for *Fmr1*^-/y^ neurons (Kolmogorov-Smirnov Sparsity: D=0.098, p=0.267; % Active pixels: D=0.100, p=0.247). Moreover, there were no differences between the distributions of genotypes for both measures on Day 1 (Kolmogorov-Smirnov Sparsity: D=0.073, p=0.613; % Active pixels: D=0.0508, p=0.942) but WT had lower values (left shift distributions) compared to *Fmr1*^-/y^ on Day 2 (Kolmogorov-Smirnov Sparsity: D=0.169, p=0.005; % Active pixels: D=0.187, p=0.001; Figure 2 D & F). All together, these data indicate that CA1 pyramidal neurons from WT rats exhibit robust experience-dependent refinement of spatial tuning between the first and second days of exploring a novel environment, but this refinement is not observed in *Fmr1*^-/y^ rats.

### Stability of CA1 pyramidal neuron firing rate maps is normal in *Fmr1*^-/y^ rats

Another key property of pyramidal neurons that has been linked to synaptic plasticity is the stability of their firing rate maps (Bett et al., 2013; Kentros et al., 1998). Interestingly, the plasticity mechanisms behind this property are molecularly dissociable from those that govern the experience-dependent increase of spatial information (Cacucci et al., 2007). To assess firing rate map stability, we calculated the correlation between the firing rate maps across consecutive sessions (yielding 5 comparisons) (Figure 3A; Figure 3 Suppl 1). This analysis only included pyramidal neurons with spatial information >0.5 bits/spike (place cells) in both sessions within a pair, to ensure that low correlations were not a result of poor spatial coding (Grieves, Wood, & Dudchenko, 2016; Roux, Hu, Eichler, Stark, & Buzsáki, 2017). Pyramidal neurons from *Fmr1*^-/y^ and WT rats had similar spatial firing rate map correlations between consecutive sessions (Figure 3 Suppl 2; LME main effect of genotype p=0.094). There was also a significant main effect of comparison (LME main effect of comparison p<0.001), driven by both groups showing lower correlations between days (Session 3 vs Session 4) than between sessions within each day (LME post-hocs p’s <0.05). The interaction between genotype and comparison was not significant (LME genotype x session comparison interaction p=0.617). Subsequent analysis of the distribution of correlations for each comparison showed that firing rate maps from *Fmr1*^-/y^ neurons were similar to those of WT neurons for all comparisons except for the comparison between the second and third session on Day 1 (Kolmogorov-Smirnov Session 1 - Session 2 WT vs *Fmr1*^-/y^: D=0.103, p=0.422; Session 2 - Session 3 WT vs *Fmr1*^-/y^: D=0.189, p<0.01; Session 3 - Session 4 WT vs *Fmr1*^-/y^: D=0.176, p=0.150; Session 4 - Session 5 WT vs *Fmr1*^-/y^: D=0.115, p=0.308; Session 5 - Session 6 WT vs *Fmr1*^-/y^: D=0.119, p=0.250) (Figure 3B).

**Figure 3.**
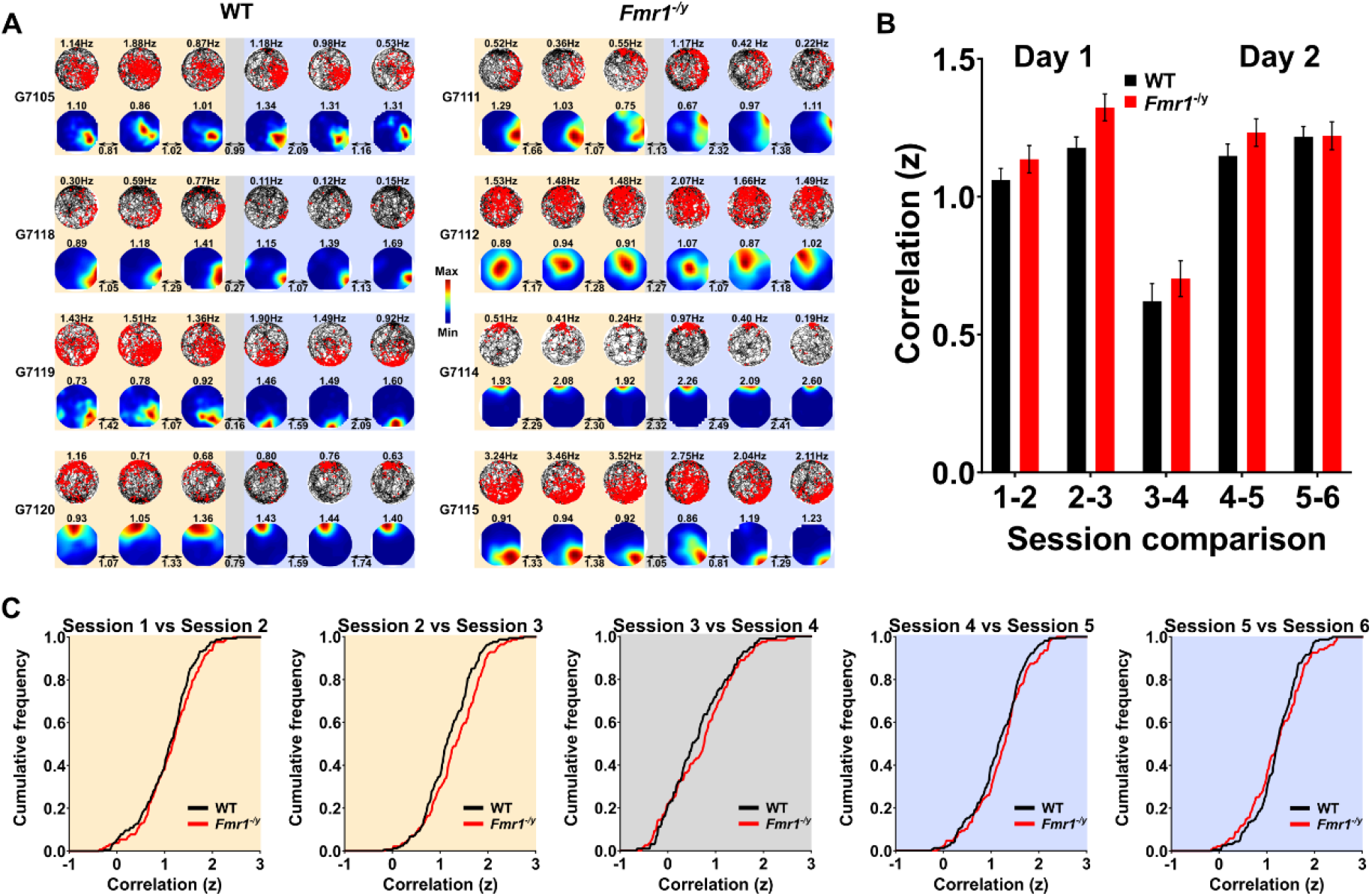
*Fmr1*^*-/y*^ CA1 place cells exhibit normal stability in their spatial discharge patterns. **(A)** Example firing rate maps from WT and *Fmr1*^*-/y*^ CA1 place cells. For each example, Top: movement trajectory (black path) and superimposed action potentials of the place cell (red dots) across the 6 sessions across the two days in the novel environment. Mean firing rate for that session is stated on top of each plot. Bottom: Smoothed firing rate maps of the same place cell, with warmer colours indicating higher rates of firing rate. Spatial information for each session is stated above the firing rate map. Correlations between the firing rate maps of successive exploration sessions (Fisher z-transformed) are stated between sessions. (**B**) Mean correlation coefficients (Fisher z-transformed) from WT and *Fmr1*^*-/y*^ rats, for all recording sessions. WT and *Fmr1*^*-/y*^ place cells exhibit similar stability of spatial discharge patterns. Data represent cell means and SEM. **(C)** Cumulative distributions of correlation coefficients (Fisher z-transformed) for all cells included in analysis (**B**) between consecutive session firing map correlations. All session comparisons, except from Session 2 vs Session 3 (*Fmr1*^*-/y*^ place cells having higher correlations) revealed the firing rate map correlation distributions were similar between WT and Fmr1^-/y^ place cells. S1vsS2: N_WT_=194, N_KO_=167, S2vsS3: N_WT_=191, N_KO_=179, S3vsS4: N_WT_=121, N_KO_=152, S4vsS5: N_WT_=190, N_KO_=161, S5vsS6: N_WT_=188, N_KO_=169. Pale yellow and pale purple backgrounds denote data from Day 1 and Day 2 respectively, while grey denotes comparison between the last session on Day 2 (Session 3) and the first session on Day 2 (Session 4).

Together, these analyses show that pyramidal neurons from *Fmr1*^-/y^ rats exhibit normal spatial stability in their firing rate maps, both between sessions within the same day and between the first two days of exposure to a novel environment. While this is consistent with stable spatial coding in *Fmr1*^-/y^ rats, this finding contrasts with the observed deficits in the experience-dependent update of spatial tuning over the two days of recordings.

### *Fmr1*^*-/y*^ CA1 pyramidal neurons are intrinsically more excitable, but receive less synaptic input from the medial entorhinal cortex

Overall, our analyses suggest that loss of FMRP leads to reduced adaptation of CA1 pyramidal neurons to a novel environment across days, maintaining high spike discharge and low spatial tuning independent of experience. A key question is whether these phenotypes result from changes in the intrinsic properties of CA1 pyramidal neurons themselves, alterations in the inputs they receive, or a combination of the two. To address this question, we performed whole-cell patch clamp recordings from adult CA1 pyramidal neurons (WT: 36 cells from 10 rats, *Fmr1*^-/y^: 48 cells from 12 rats) from the dorsal hippocampus matched to the location of the *in vivo* recordings. First, we measured the intrinsic excitability of CA1 pyramidal neurons in response to depolarizing current steps (0 - 400 pA, 25 pA steps, 500 ms duration), which in WT neurons resulted in repetitive action potential discharge (Figure 4A). The same stimuli delivered to CA1 pyramidal neurons from *Fmr1*^-/y^ rats resulted in consistently increased AP discharge over the range of currents tested (Figure 4B). Comparison of the slopes of individual action potential discharge to increasing current revealed an increase in the overall slope in *Fmr1*^-/y^ CA1 pyramidal neurons, compared to WT (Figure 4C; LME: p=0.04). We have recently shown in the CA1 of juvenile *Fmr1*^-/y^ mice that such changes in intrinsic cell excitability are correlated with altered membrane potential, reduced threshold to fire, and longer axon initial segments (AIS) (Booker et al., 2020). Comparing CA1 pyramidal neurons between WT and *Fmr1*^-/y^ rats, we observed no difference in passive properties such as resting membrane potential (Figure 4D; LME: p=0.45), input resistance (Figure 4E; LME: p=0.51), rheobase current (Figure 4F, *p*=0.18) or action potential threshold (Figure 4G; LME: p=0.46). However, the medium after-hyperpolarisation (mAHP) was reduced by 13% in *Fmr1*^-/y^ CA1 pyramidal neurons compared to WT (Figure 4H, p=0.05). Consistent with the similar action potential thresholds measured between genotypes, we observed no difference in the AIS length, as assessed by AnkyrinG immunostaining (Figure 4I), between *Fmr1*^-/y^ and WT rats (Figure 4I; LME: p=0.33).

**Figure 4.**
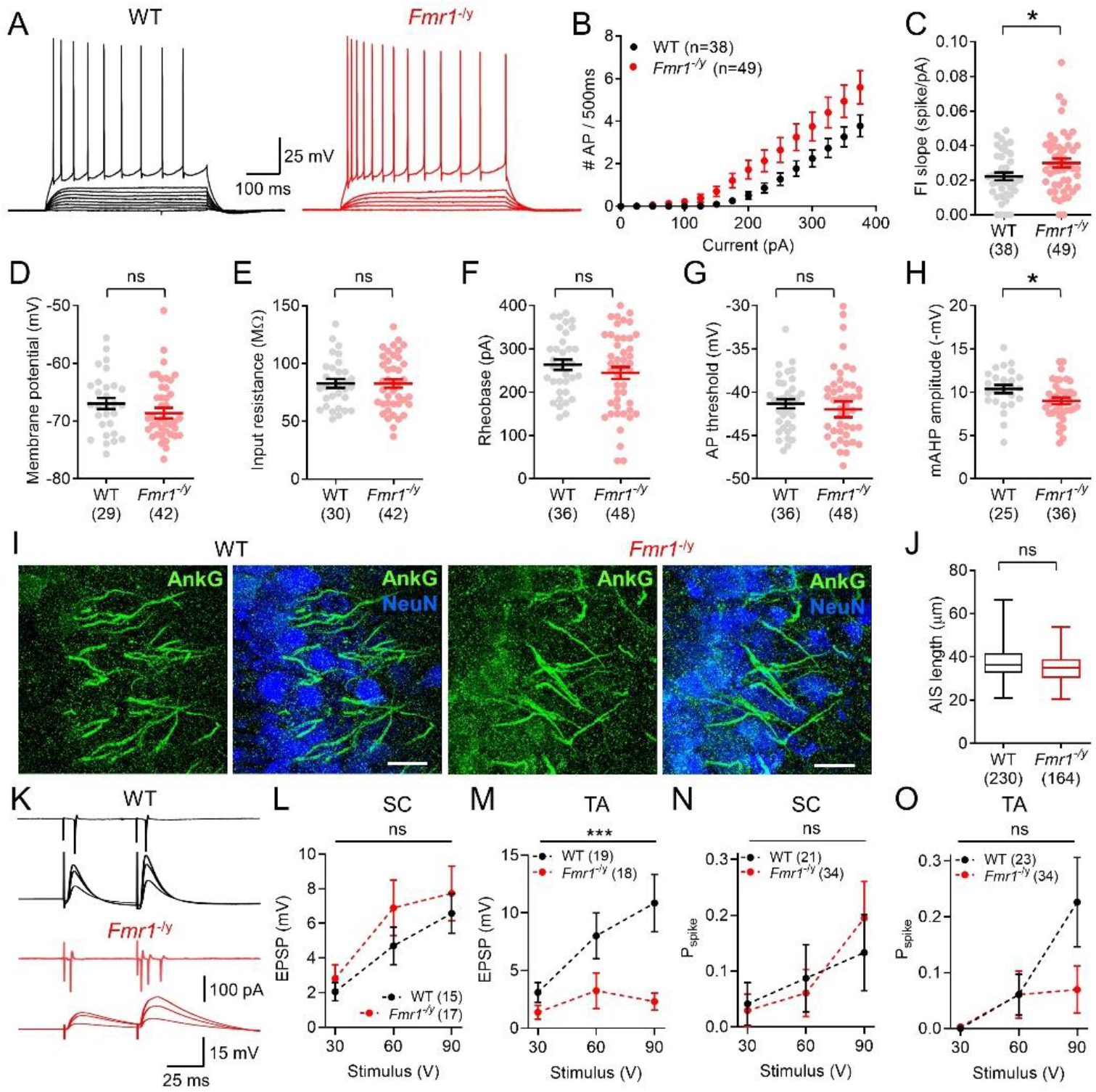
CA1 pyramidal neurons in slices from *Fmr1*^*-/y*^ rats display increased excitability, which is correlated with reduced synaptic inputs from the medial entorhinal cortex. **A** Representative traces of recordings from CA1 pyramidal neurons in the dorsal hippocampus from WT (black) and *Fmr1*^*-/y*^ (red) rats, in response to depolarizing current injections (0 - 400 pA, 25 pA steps, 500 ms duration). **B** Plot of the action potential discharge in 500 ms compared to injected current for all recorded CA1 pyramidal neurons in WT (black) and *Fmr1*^*-/y*^ (red) rats. Data as shown as the mean response recorded per rat, with number of rats indicated. As measured from the zero-current potential, average resting membrane potential (**C**) and input resistance (**D**) are quantified. Data from individual neurons are shown overlain as filled circles, and the number of tested neurons shown below in parentheses. Average data are plotted for the rheobase current (**E**) and the voltage threshold (**F**) and medium afterhyperpolarisation (mAHP) amplitude (**G**), the latter two measured from the first action potential at rheobase. **H** Visualization of the AIS in flattened confocal z-stacks as revealed by immunofluorescent labelling for AnkyrinG (AnkG, green pseudocolour) and merged with NeuN (blue pseudocolour) from WT (left) and *Fmr1*^*-/y*^ rats (right). Scale bars shown: 20 µm. (**I)** Quantification of AIS length, displayed as the average of individual animals. **J** Representative traces from cell attached recordings (upper traces) and whole cell recordings (lower traces) following stimulation of str. lacunosum moleculare to distal CA1, from WT (top) and *Fmr1*^*-/y*^ rats (bottom). EPSP data is shown as the average traces in response to 30 V, 60 V, and 90 V stimulation intensities. Average data for EPSP amplitude in response to the same stimulation intensities, delivered to the Schaffer-Collateral (SC, **K**) and temporoammonic (TA, **L**) pathways. Average spike probability, as measured in cell-attached recordings, are shown for SC (M) and TA (N) paths. For **K-N**, all graphs display the result from 2-way ANOVA for genotype shown above the chart and number of tested neurons indicated in parentheses. All data is shown as mean ± SEM. Statistics shown as: ns - p > 0.05, * - p < 0.05, and *** - p < 0.001 from GLMM analysis, except panels B, K, L, M, N which are the result of 2-way ANOVA for genotype.

We next examined the synaptic strength of the two major excitatory inputs to CA1: the Schaffer-Collateral (SC) inputs from CA3 neurons in *stratum radiatum* and *oriens* and the temporoammonic (TA) inputs from layer 3 of the medial entorhinal cortex (MEC3) onto the distal dendrites in *stratum lacunosum-moleculare* (Amaral and Witter, 1989). We have previously shown reduced strength of TA synapses with CA1 pyramidal neurons in the mouse model of FXS (Booker et al., 2020). Cell attached and whole cell recordings from CA1 pyramidal neurons were performed in the presence of 50μM picrotoxin in order to block inhibitory neurotransmission (Figure 4K and Figure 4 Suppl 1A&B). In whole-cell patch clamp recordings, stimulation of SC afferents in *stratum radiatum* did not show a difference between genotypes in the resulting excitatory postsynaptic potential (EPSP) measured over the range of stimulation intensities tested (30, 60 and 90 V; Figure 4L; 2-way ANOVA main effect of genotype F_(1,68)_=2.03, *p*=0.16). In contrast, there was a robust decrease in the recruitment of EPSPs in CA1 pyramidal neurons from the *Fmr1*^-/y^ rats in response to TA stimulation compared to WT rats (Figure 4M; 2-way ANOVA: main effect of genotype F_(1,98)_=15.4, p=0.0002). This was reflected by a similar input-output slope for SC stimulation between genotypes, but a reduced slope for TA inputs in the *Fmr1*^-/y^ rats (Figure 4 Suppl 1C, K-W_(4,50)_=16.12, p=0.001 Kruskal-Wallis test). We also observed bidirectional effects on short-term plasticity in the different inputs. Specifically, at the SC inputs paired-pulse ratio (PPR) was lower in slices from *Fmr1*^*-/y*^ than from those of WT rats at all stimulation intensities (Figure 4 Suppl 1D; 2-way ANOVA: main effect of genotype F_(1,54)_=10.6, p=0.002), whereas at the TA inputs PPR was higher in the *Fmr1*^*-/y*^ rats at the 90 V stimulation intensity (Figure 4 Supple 1E; 2-way ANOVA genotype x stimulus interaction F_(2,85)_=3.92, *p*=0.024, 90 V: t_(13,14)_=2.47, p=0.05). These reductions in synaptic input were not associated with gross anatomical changes in CA1 pyramidal neurons (Figure 4 Suppl 2A), as Sholl analysis of recorded neurons displayed no difference in branching pattern (Figure 4 Suppl 2A). Furthermore, there was no difference in total or compartment specific dendritic lengths between genotypes (Figure 4 Suppl 2C-F). Finally, there was a tendency towards reduced dendritic protrusions in the distal apical dendrites, consistent with a reduction in synaptic number at TA inputs, which was not apparent at dendrites aligned to SC afferents (Figure 4 Suppl. 3).

To determine whether the increased AP discharge observed in CA1 pyramidal neurons is sufficient to overcome the reduced synaptic strength observed in response to electrical stimulation of the TA pathway, we next asked if spike output was altered. In cell-attached recordings from pyramidal neurons under the same conditions, we performed paired-pulse stimulation at SC and TA synapses to CA1. We observed a similar recruitment of CA1 pyramidal neurons to SC pathways stimulation in *Fmr1*^*-/y*^ and WTs, both in terms of spike probability (Figure 4N; 2-way ANOVA: main effect of genotype F_(1,156)_=0.032, *p*=0.86) and in terms of number of cells that produced a spike in any response to any stimulus (Figure 4 Suppl 1F; χ^2^=0.30, *p*=0.59). In contrast, the overall spike probability following TA afferent stimulation was not different between genotypes at 30 V and 60 V stimulation (LME: *p*=0.99), but tended towards lower spiking in *Fmr1*^*-/y*^ than in WT rats at 90 V (LME: p=0.06), suggesting a level of compensation, at least at lower stimulation intensities (Figure 4O; 2-way ANOVA: main effect of genotype F_(1,163)_=2.276, *p*=0.13). However, the overall recruitment of individual CA1 pyramidal neurons was reduced by approximately 3-fold (Figure 4 Suppl 1F; χ^2^=4.37, *p*=0.04), indicating a degree of information loss. Despite the reduction in CA1 pyramidal neuron recruitment, when present, spikes showed a similar coefficient of variation of spike times between genotypes, in the absence of GABA_A_ receptor mediated inhibition (Figure 4 Suppl 1G; K-W_(4,44)_=0.23, *p*=0.97).

Together, these data show that CA1 pyramidal neurons in the dorsal hippocampus of the *Fmr1*^-/y^ rat are hyperexcitable, which results from a decrease in medium AHP amplitude. This hyperexcitability is sufficient to overcome the reduced synaptic strength following TA stimulation received by *Fmr1*^-/y^ CA1 p-yramidal neurons.

### Pyramidal neurons from *Fmr1*^-/y^ rats exhibit decreased phase modulation by gamma oscillations

The firing patterns of CA1 pyramidal neurons can be temporally organized by both gamma and theta oscillatory frequencies of local field potentials (LFP) while an animal explores its environment (Fernández-Ruiz et al., 2017; Lasztóczi & Klausberger, 2016). The different frequency bands reflect hippocampal circuit organization as well as unique inputs (Colgin, 2016; Lisman & Jensen, 2013) (Figure 5 Suppl 1A). Analysis of LFP oscillatory power during mobility revealed no significant differences between genotypes or across days in the mean power of theta, slow gamma or medium gamma during these epochs (Figure 5A-C; Figure 5 Suppl 1B-E; Theta: RM ANOVA: genotype F_(1,12)_=0.880, p=0.367, day F_(1,12)_=0.322, p=0.581, genotype x day F_(1,12)_=0.666, p=0.430; Slow gamma: RM ANOVA: genotype F_(1,12)_=1.988, p=0.184, day F_(1,12)_=0.120, p=0.735, genotype x day F_(1,12)_=1.613, p=0.228; Medium gamma: RM ANOVA: genotype F_(1,12)_=3.109, p=0.103, day F_(1,12)_=0.437, p=0.521, genotype x day F_(1,12)_=3.343, p=0.092). For theta, there was a significant decrease in power across sessions within each day (session F_(2,24)_=4.580, p=0.021) but this did not differ between genotypes (session x genotype interaction F_(2,24)_=0.337, p=0.717). Neither slow nor medium gamma power differed across sessions within each day (p’s>0.05).

**Figure 5.**
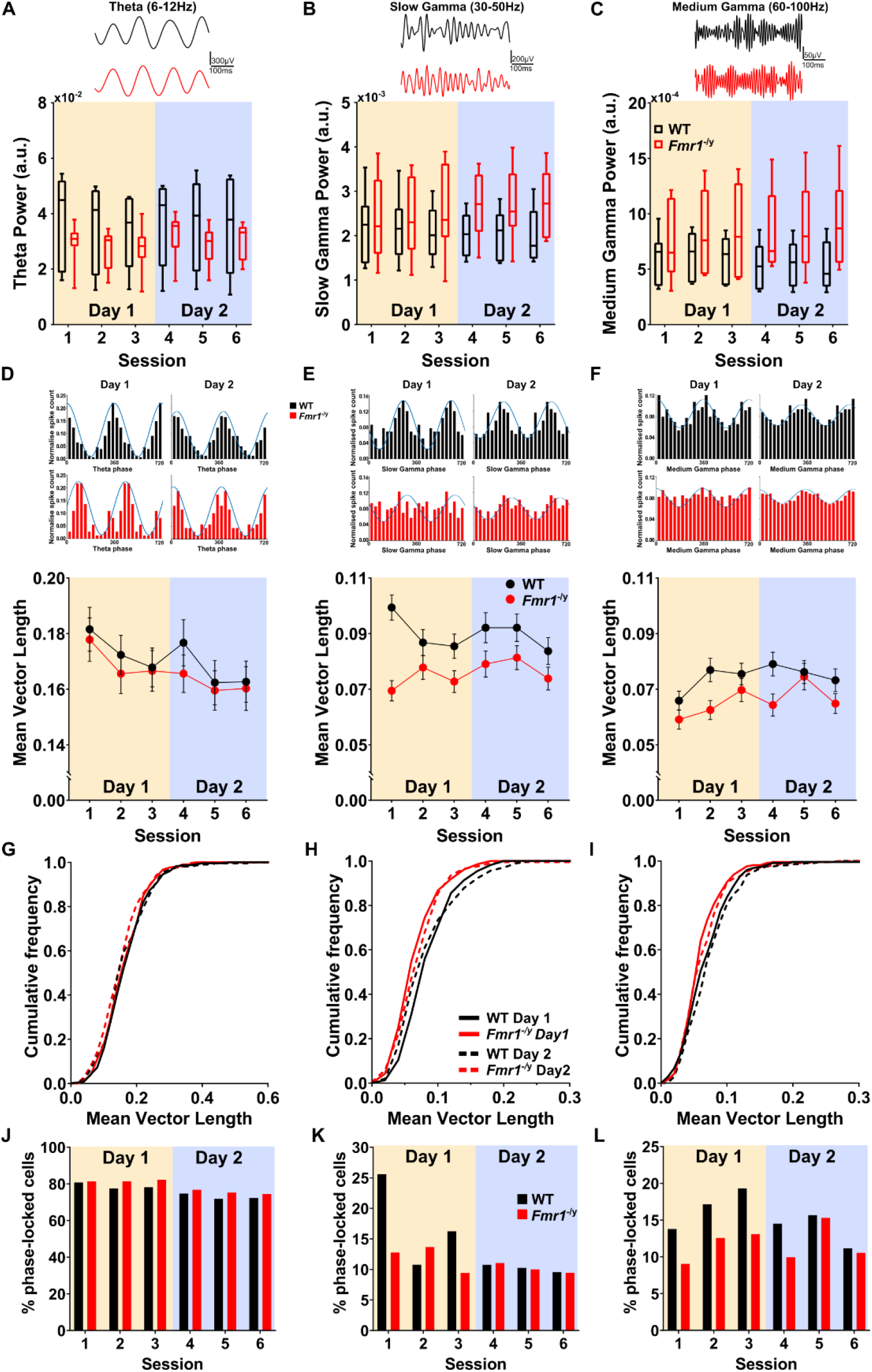
*Fmr1*^*-/y*^ CA1 pyramidal neurons exhibit decreased phase modulation by gamma oscillations. (**A**) Box plots depicting LFP power in the theta (6-12 Hz) for each recording session. Top: Example CA1 LFP traces bandpass filtered for theta from a WT (black) and an *Fmr1*^*-/y*^ (red) rat. The middle line represents rat median, upper and lower end of the box represents 95^th^ and 5^th^ percentile, error bars represent maximum and minimum values. The same for slow gamma (**B**) and medium gamma (**C**). N_WT_ = 7, N_KO_ =7. (**D**) Average mean vector length (MVL), quantifying the strength of phase locking to theta oscillations across six recording sessions. Top: Representative firing phase distribution of a single principal neuron along a theta cycle. Blue, curve fitting with von Mises distribution. Same for slow gamma oscillations (**E**) and medium gamma oscillations (**F**). Data represent cell means and SEMs. (**G**) Cumulative distribution of MVL for theta for unique pyramidal neurons recorded from WT and *Fmr1*^*-/y*^ rats during three sessions of Day 1 and three sessions of Day 2. For neurons identified in more than one session within a day, the mean MVL across these sessions is included in the analysis. No differences in the distributions of MVL between WT and *Fmr1*^*-/y*^ pyramidal neurons. (**H**) Same as (**G**) for slow gamma oscillations. Distribution of *Fmr1*^*-/y*^ pyramidal neurons is significantly shifted towards lower MVL values compared to that of WT pyramidal neurons in both days of recording. (**I**) Same as (**G**) medium gamma oscillations. Distribution of *Fmr1*^*-/y*^ pyramidal neurons is significantly shifted towards lower MVL values compared to that of WT pyramidal neurons in both days of recording. (**J**) The proportion of significantly phase-locked (Rayleigh p<0.05) pyramidal neurons to theta oscillations across six recording sessions. No differences between genotypes were observed. (**K**) Same as (**J**) for slow gamma oscillations. A higher proportion of WT pyramidal neurons is significantly modulated by slow gamma oscillations on the first recording session compared to all other recording session. This phenomenon is not evident in *Fmr1*^*-/y*^ neurons. (**L**) Same as (**J**) for medium gamma oscillations. No differences between genotypes were observed. N_WT-D1_=222, N_WT-D2_=207, N_KO-D1_=211, N_KO-D2_=205. Pale yellow and pale purple backgrounds denote data from Day 1 and Day 2 respectively.

To examine the temporal organization of CA1 pyramidal neurons firing relative to theta, slow gamma and medium gamma oscillations, we computed the mean vector length (MVL) of the phase distribution of spikes for each pyramidal neuron for each of these frequency bands. This provides a measure of how consistently a neuron fires at a specific phase of the oscillation. We first compared the MVLs of all recorded CA1 pyramidal neurons in *Fmr1*^-/y^ and WT rats across sessions and days.

The MVL for theta did not differ between genotypes or between days according to our LME analyses (Figure 5D; Figure 5 Suppl 2A&B; Figure 5 Suppl 3; main effect genotype p>0.05, main effect of day p>0.05). However, there was a significant decrease in MVL for both genotypes across sessions within each day (session-in-day effect p=0.026). Similarly, the distribution of MVL values did not differ between genotypes or days (Figure 5G; Kolmogorov-Smirnov Day 1 WT vs *Fmr1*^-/y^: D=0.071, p=0.649; Day 2 WT vs *Fmr1*^-/y^: D=0.095, p=0.309; WT Day1 vs Day2: D=0.095, p=0.291; *Fmr1*^-/y^ Day1 vs Day2: D=0.098, p=0.272).

In contrast, neurons from *Fmr1*^-/y^ rats showed significantly less phase locking (shorter MVLs) to slow gamma than those from WT rats, but no differences between days were found (Figure 5E; Figure 5 Suppl 2C&D; Figure 5 Suppl 3; LME: main effect of genotype p=0.022; main effect of day p=0.538; genotype x day interaction p=0.109). Analysis of MVL distribution corroborated the LME analysis results showing a reduction in spiking modulation by slow gamma for *Fmr1*^-/y^ rats on both days (Figure 5H; Kolmogorov-Smirnov Day 1 WT vs *Fmr1*^-/y^: D =0.273, p <0.001; Day 2 WT vs *Fmr1*^-/y^: D=0.154, p=0.015; WT Day1 vs Day2: D=0.132, p=0.049; *Fmr1*^-/y^ Day1 vs Day2: D=0.092, p=0.348).

Our analyses for medium gamma yielded mixed results: the LME model approach did not show a significant effect of genotype, day or genotype x day interaction (Figure 5F; Figure 5 Suppl 2E&F; Figure 5 Suppl 3; LME: main effect of genotype p=0.136; main effect of day p=0.082; genotype x day interaction p=0.935). However, analysis of the distribution of MVLs indicated less phase locking to medium gamma in neurons from *Fmr1*^-/y^ than WT rats on both days with no changes over days (Figure 5I; Kolmogorov-Smirnov Day 1 WT vs *Fmr1*^-/y^: D=0.169, p=0.004; Day 2 WT vs *Fmr1*^-/y^: D=0.174, p=0.004; WT Day1 vs Day2: D=0.094, p=0.306; *Fmr1*^-/y^ Day1 vs Day2: D=0.099, p=0.259).

We further quantified the proportion of recorded CA1 pyramidal neurons that were significantly phase locked to theta, slow gamma and medium gamma oscillations (MVL Rayleigh p<0.05). In line with previous analyses, we detected no significant difference between WT and *Fmr1*^-/y^ in the proportion of neurons (across all rats of each genotype) that showed significant phase locking to theta. However, we found a significant effect of session (Figure 5J; Figure 5 Suppl 2 G; Log-likelihood ratio: session G^2^=12.50, p=0.029; genotype G^2^=12.50, p=0.129; for details see methods).In contrast to theta, the proportion of *Fmr1*^-/y^ CA1 neurons that were significantly phase locked to slow gamma was not modulated by experience, the same as neurons from WT rats. We found a significant genotype x session interaction (Figure 5K; Figure 5 Suppl 2 H; Akaike’s Criterion probability ratio = 1.515, AICc =0.831; for details see methods). Further comparison between genotypes during each session revealed that the proportion of slow gamma phase locked neurons was higher in WT that rats in Sessions 1 and 3 of Day 1, but not for any other session (two proportion z-test, WT vs *Fmr1*^-/y^ Session 1: p<0.001; Session 3: p=0.023). Moreover, a significantly higher proportion of WT neurons were phase-locked in Session 1 than any other session, whereas there were no differences across sessions for *Fmr1*^-/y^ rats (two proportion z-test WT, Session 1 vs Session 2 p<0.001; Session 1 vs Session 3 p=0.014; Session 1 vs Session 4 p<0.001; Session 1 vs Session 5 p<0.001; Session 1 vs Session 6 p<0.001; KO all comparisons p>0.05). Finally, we found a significant difference between WT and *Fmr1*^-/y^ rats in the proportion of pyramidal neurons that were phase-locked to medium gamma(Figure 5L; Figure 5 Suppl 2 I; Log-likelihood ratio: Session G^2^=8.42, p=0.135; Genotype G^2^=6.24, p=0.013; for details see methods). This finding corroborated the MVL distribution analysis (Figure 5I).

Collectively, these analyses reveal that CA1 pyramidal neurons from *Fmr1*^-/y^ rats are less modulated by gamma oscillations compared to neurons in WT rats, and, with some measures. These findings indicate that the timing of CA1 pyramidal neuron firing relative to gamma oscillations, particularly slow gamma, is less consistent in *Fmr1*^-/y^ compared to WT rats. In addition, in WT (but not in *Fmr1*^-/y^ rats) significantly more neurons were phase-locked to slow gamma in Session 1 on Day 1 (when the environment was completely novel) than in any other session. This provides further evidence of decreased experience-dependent changes in CA1 neuronal activity in *Fmr1*^-/y^ rats.

### CA1 pyramidal neurons from *Fmr1*^-/y^ rats discharge preferentially during the ascending phase of theta

Another measure that can be used to characterize the organization of ensemble activity patterns is the timing of CA1 pyramidal neuron spiking relative to the theta and gamma oscillations. Inputs to the CA1 area of hippocampus have specific temporal organization, with inputs from CA3 arriving predominantly during the descending phase of CA1 theta, and inputs from MEC3 neurons primarily during the ascending phase and peak of theta (Mizuseki, Sirota, Pastalkova, & Buzsáki, 2009; Schomburg et al., 2014) (Figure 6A). Moreover, CA1 pyramidal neurons change their preferred theta phase (to later phase of the theta cycle) in response to novelty (Lever et al., 2010). Less is known about the functional significance of CA1 pyramidal neuron firing during specific gamma phases, although a correlation between slow gamma phase preference of CA1 pyramidal neurons and the position of their place fields in a linear track has been reported (Zheng, Bieri, Hsiao, & Colgin, 2016). Taking into account that *Fmr1*^-/y^ rats exhibit decreased phase modulation by slow gamma oscillations and the fact that inputs from CA3 (reflected in slow gamma power) arrive predominantly during the descending phase of theta, we predicted that *Fmr1*^-/y^ CA1 pyramidal neurons would show decreased firing preference for the descending phase of theta compared to WT neurons.

**Figure 6.**
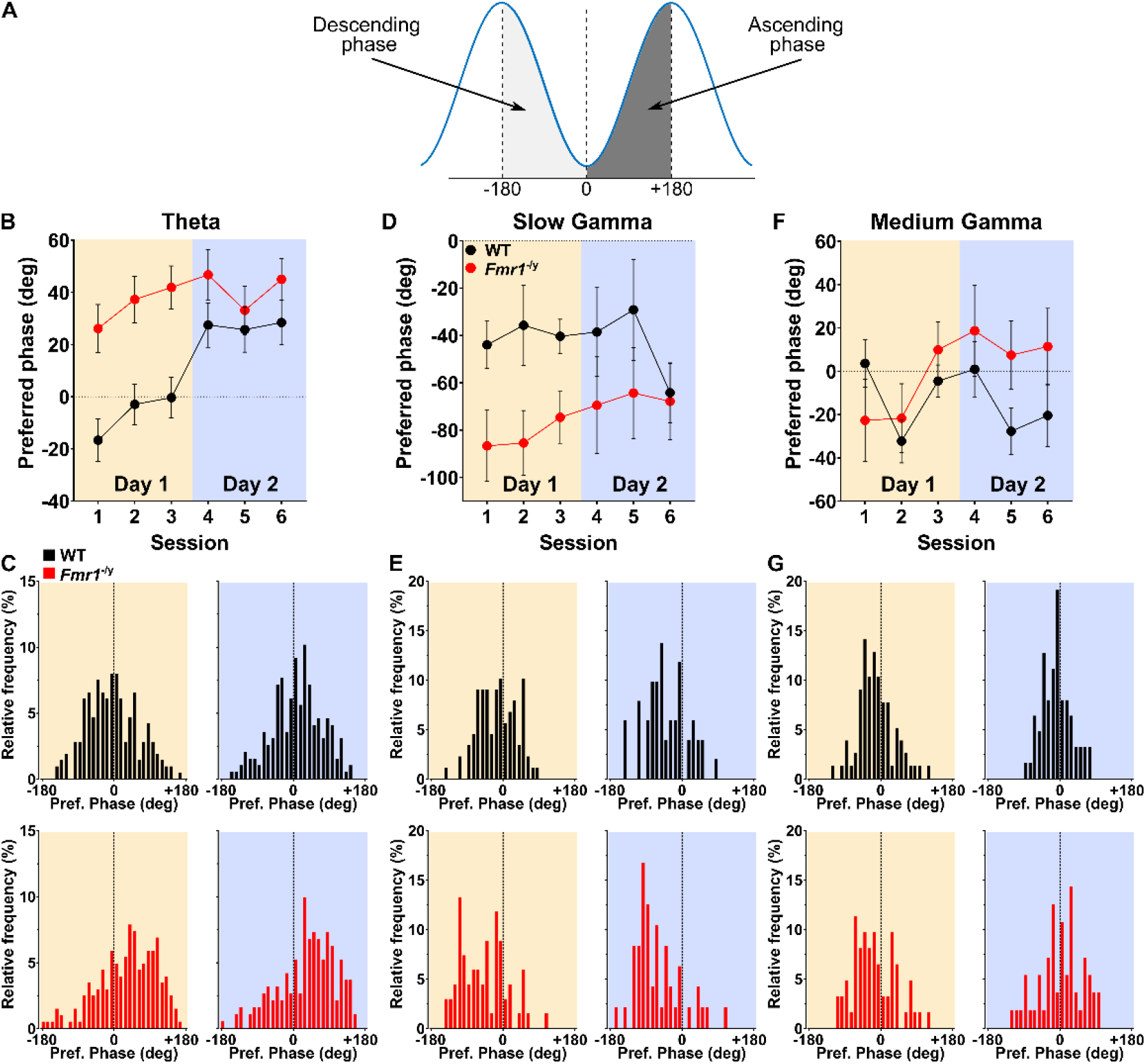
*Fmr1*^*-/y*^ CA1 pyramidal neurons discharge later during theta cycle and do not show experience-dependent changes in theta phase preference. (**A**) Schematic depiction of oscillation phases. The oscillation troughs were defined as 0°. (**B**) Mean preferred theta discharge phase for significantly phase-locked (Rayleigh p<0.05) pyramidal neurons during the six recording sessions. Same for slow gamma oscillations (**D**) and medium gamma oscillations (**F**). (**C**) Distribution of preferred theta discharge phase for unique pyramidal neurons recorded from WT (Top-black) and *Fmr1*^*-/y*^ (Bottom-red) rats during Day 1 (left) and Day 2 (right). For neurons identified in more than one session within a day, the mean preferred theta discharge phase across these sessions was included in the distribution. Same for slow gamma oscillations (**E**) and medium gamma oscillations (**G**). Theta: N_WT-D1_=214, N_WT-D2_=197, N_KO-D1_=199, N_KO-D2_=187, SGamma: N_WT-D1_=89, N_WT-D2_=51, N_KO-D1_=65, N_KO-D2_=47, SGamma: N_WT-D1_=78, N_WT-D2_=63, N_KO-D1_=60, N_KO-D2_=56. Pale yellow and pale purple backgrounds denote data from Day 1 and Day 2 respectively.

We analysed the preferred theta phase as well as the preferred slow and medium gamma phases of CA1 pyramidal neurons from *Fmr1*^-/y^ and WT rats. Only neurons with significant phase modulation (based on MVL Rayleigh p<0.05) were included in our analysis. Due to the circular nature of the data, we used the Harrison-Kanji two-factor test for circular data and the Watson-Williams multi-sample test to explore differences between genotypes in this dataset. We found that WT pyramidal neurons discharged preferentially around the trough of theta during the first day of exposure to the novel environment, but more towards the ascending phase on the second day. In *Fmr1*^-/y^ rats, pyramidal neurons fired predominantly during the ascending phase of theta on Day 1 and Day 2 (Harrison-Kanji test, main effect of genotype p<0.001; genotype x day interaction p=0.0025; Watson-Williams test, WT, Day1 vs Day2 p<0.001; *Fmr1*^-/y^, Day1 vs Day2 p=0.858; Day1, WT vs *Fmr1*^-/y^ p<0.001; Day2, WT vs *Fmr1*^-/y^ p<0.001) (Figure 6B&C). Analysis of the slow gamma preferred spiking phase revealed that WT and *Fmr1*^-/y^ pyramidal neurons discharge primarily during the descending phase of the oscillation on both Day 1 and Day 2 (Harrison-Kanji test, main effect of day: p=0.614) (Figure 6D&E). However, *Fmr1*^-/y^ neurons discharged at an earlier phase of slow gamma than WT neurons (Harrison-Kanji test, main effect of genotype: p=0.003; genotype x day interaction: p=0.664). Finally, medium gamma discharge phase preference was not significantly different between WT and *Fmr1*^-/y^ rats (Harrison-Kanji test, main effect of genotype: p=0.056; genotype x day interaction: p=0.219) (Figure 6F&G).

Together, these analyses revealed that *Fmr1*^-/y^ pyramidal neurons exhibit differences in their preferred phase discharge compared to their WT counterparts. For theta, this difference is more prominent during the exploration of a novel environment (Day 1), while for slow gamma, the difference persists throughout both days of recordings.

## Discussion

Individuals with FXS have intellectual disability and ASD, two features in which hippocampal pathophysiology has been implicated (Barnes et al., 2015; Cooper et al., 2017; Koenig et al., 2020; Till et al., 2015; Witton et al., 2015). While much is known about the cellular physiology and biochemical changes associated with the loss of FMRP, less is known about how cellular pathology leads to circuit dysfunction. The recent findings in the mouse models of FXS (Arbab et al., 2018; Arbab, Pennartz, & Battaglia, 2018; Boone et al., 2018; Dvorak et al., 2018; Radwan, Dvorak, & Fenton, 2016; Talbot et al., 2018) and the conserved hippocampal pathophysiology between *Fmr1*^-/y^ mice and rats (Till et al., 2015) led us to predict that aspects of temporal coordination of both spiking activity and LFP would also be affected in *Fmr1*^-/y^ rats. In the current study, using *in-vivo* electrophysiological recordings, we show that CA1 pyramidal neurons in the *Fmr1*^*-/y*^ rat model of Fragile X syndrome display significant impairments in appropriate adaptation to repeated presentations of a novel spatial environment. Specifically, we show that while WT neurons exhibit experience-dependent changes in firing across days in a novel environment, with commensurate changes in spike bursting, spatial coding, and the timing of spikes relative to the local field potential, these changes are absent in *Fmr1*^*-/y*^ rats. Our findings suggest that the *Fmr1*^*-/y*^ rats may persistently experience environments as novel, despite repeated exposure across days. This deficient habituation to novelty is a core clinical feature of FXS (Bruno et al., 2014; Ethridge et al., 2016). Our *in vivo* findings are supported by *ex vivo* recordings that provide evidence for single cell hyperexcitability in the face of reduced synaptic inputs from the medial entorhinal cortex. Using *ex-vivo* electrophysiological recordings, we showed that the circuit function abnormalities we observe may be, at least in part, the result of reduced synaptic inputs from MEC3 to CA1. Together, these data provide novel insight into how the absence of FMRP may result in altered adaptation to novel spaces.

### Hippocampal CA1 pyramidal neuron activity does not exhibit experience-dependent decrease in Fmr1^-/y^ rats

On the first day in a novel environment, we found that key features of CA1 pyramidal neuron firing, including rate and burstiness, spatial information coding and sparsity, did not differ between WT and *Fmr1*^-/y^ rats. However, when re-exposed to the same environment the next day, WT rats displayed reduced firing and burstiness, with increased spatial selectivity, as previously described in CA1 pyramidal neurons in WT rats (Brandon, Koenig, Leutgeb, & Leutgeb, 2014; Karlsson & Frank, 2008; Nitz & McNaughton, 2004), but these changes were absent in *Fmr1*^*-/y*^ rats. The decrease in mean firing rate in WT rats is thought to be mediated by synaptic plasticity, whereby synaptic inputs to CA1 pyramidal neurons are stronger in a novel environment than in a familiar environment (Cohen, Bolstad, & Lee, 2017). Strong synaptic inputs during exploration of the novel environment (Day 1) would therefore lead to higher spiking activity in WT CA1 pyramidal neurons than on the second day of recording in the same environment, when inputs are weaker. We propose that the increased excitability of *Fmr1*^-/y^ CA1 pyramidal neurons is able to compensate for the diminished synaptic inputs from MEC3 to CA1 in the *Fmr1*^-/y^ rats(Figure 4), allowing the CA1 neurons in *Fmr1*^-/y^ rats to show similar (high) levels of activity in the novel environment as WT rats on Day 1. However, the increased excitability may lead to a ceiling effect in the activity of *Fmr1*^*-/y*^ CA1 pyramidal neurons, making discrimination between novel (higher MEC3 to CA1 inputs) and familiar (lower MEC3 to CA1 inputs) environments suboptimal. We have previously shown a similar functional effect (i.e. increased CA1 pyramidal neuron excitability) in the dorsal hippocampus of a mouse model of FXS (Booker et al., 2020). Interestingly, the mechanisms mediating increased CA1 excitability appears to differ between rats and mice (e.g. increased AIS length), suggesting that while this alteration to hippocampal circuit function in the *Fmr1*^-/y^ rodents may be species independent, the mechanism by which mice and rats compensate for such change is divergent. Such compensation of synaptic deficits by individual neurons may in fact lead to many features of FXS and ASD/ID more generally (Booker & Kind, 2021).

In addition to a decrease in mean firing rate, WT pyramidal neurons also showed decreased bursting probability on Day 2 compared to Day 1, consistent with firing rate correlating to burst probability (Mizuseki & Buzsáki, 2013). This phenomenon was absent in *Fmr1*^-/y^ pyramidal neurons. Given that the likelihood of bursting depends largely on the intrinsic excitability of neurons (Jarsky et al., 2008; Harris et al., 2001), the pattern in *Fmr1*^-/y^ neurons may also be explained by the homeostatic increase in the excitability of *Fmr1*^*-/y*^ CA1 pyramidal neurons (Figure 4). Specifically, reduction of mAHP has been shown to cause increased bursting (Gu et al., 2005). In our data, the increase in single cell excitability was unrelated to a change in the length of the axon initial segment, which differs from our previous findings in *Fmr1*^-/y^ mice (Booker et al., 2020). The reduced mAHP we observed may explain the hyperexcitability in CA1 pyramidal cells from adult *Fmr1*^-/y^ rats, which is consistent with previous findings in the ventral hippocampus in mice (Ordemann et al, 2021).

Previous *in-vivo* studies in *Fmr1*^-/y^ mice have reported inconsistent firing rate phenotypes in CA1 pyramidal neurons. Consistent with our finding of higher firing in *Fmr1*^-/y^ rats on the second day of recording, CA1 pyramidal neurons of *Fmr1*^-/y^ mice were found to exhibit higher firing rates than WT mice in a very familiar environment (Boone et al., 2018). However, other studies have reported no differences in firing rate between *Fmr1*^-/y^ and WT mice exploring familiar environments (Talbot et al., 2018), or over 2 days in a novel environment (Arbab et al., 2018). While the result of Arbab et al. (2018) in mice appears to be inconsistent with our finding in rats, it is notable that Arbab et al. did not observe experience-dependent changes in firing rate between days even in WT mice (as also reported in a previous report in WT mice (Cacucci et al., 2007). Therefore, the failure to detect differences between WT and *Fmr1*^-/y^ mice in their study is perhaps not surprising and may indicate some species divergence in hippocampal adaptation to environmental novelty. Indeed, mice need a substantially longer period of familiarization and training to perform certain tasks compared to rats (Colacicco et al., 2002; Jaramillo and Zador, 2014).

### Refinement of spatial information coding in the CA1 region of the hippocampus is impaired in Fmr1^-/y^ rats

The spatial information conveyed by hippocampal pyramidal neurons has previously been shown to increase sharply between the first and second days of exploration in a novel environment in mice (Cacucci et al., 2007) and in rats learning a set of spatial rules in an open field (Retailleau & Morris, 2018). Our findings show a similar experience-dependent increase in spatial information of CA1 pyramidal neurons in WT rats between the first two days of exploration of a novel environment, but that this is absent in *Fmr1*^-/y^ rats. This difference between WT and *Fmr1*^-/y^ rats may be due, at least in part, to the decreased strength of synaptic inputs to CA1 from MEC3 that we observed in *Fmr1*^-/y^ rats in our *ex-vivo* experiments, and which we have previously reported in the *Fmr1*^-/y^ mouse (Booker et al., 2020). Lesions of MEC3 result in large reductions in the spatial information of CA1 pyramidal neurons in rats exploring a familiar environment (where control animals show high spatial information), whereas they have much more subtle effects on spatial information in a novel environment (in which spatial information is low in both lesioned and control rats) (Brun et al., 2008). This indicates that in WT rats, MEC3 inputs to CA1 typically contribute to the experience-dependent refinement of the spatial coding, and that the failure of *Fmr1*^-/y^ pyramidal neurons to show an increase in spatial information upon familiarization to the novel environment across days may result from the reduced strength of these synaptic inputs.

While spatial coding showed the expected refinement between days 1 and 2 in WT rats, the between-session stability of firing rate maps of pyramidal neurons did not change from Day 1 to Day 2 in either WT or *Fmr1*^-/y^ rats. Contrary to our predictions, the firing rate maps of *Fmr1*^-/y^ rats were as stable as those of WT rats, both within and across days. This was unexpected, as previous findings in *Fmr1*^-/y^ mice indicated lower firing rate map stability across repeated exploration of a novel environment (Arbab et al., 2018). That being said, in our study both the daily session number (3) session duration (10 min) and within-day inter-trial intervals (10 min) differ from those of Arbab et al. (2018) (two 30-min sessions on Day 1 with a 2 h inter-trial interval), which could have affected firing rate map stability both within and across days.

### Hippocampal LFP power is largely unaffected by the loss of FMRP

Our LFP recordings from the CA1 pyramidal neuron layer did not reveal any significant differences between genotypes, or any experience-dependent changes in the power of theta (6-12 Hz), low-range gamma (30-50 Hz) or mid-range gamma (60-100 Hz) band activity. At face value, these findings contradict previous reports in *Fmr1*^-/y^ mice, and may suggest that our experiment was underpowered. However, the non-significant trends towards higher gamma power in *Fmr1*^-/y^ rats, particularly on the second day of exploration of the novel environment, echo the findings from a number of studies reporting that gamma power is stronger in the hippocampus of *Fmr1*^-/y^ mice (Arbab et al., 2018; Boone et al., 2018; Dvorak et al., 2018). Beyond the hippocampus, increased gamma-band power has been reported in the frontal and temporal cortex of *Fmr1*^-/y^ mice (Jonak, Lovelace, Ethell, Razak, & Binder, 2020), the parietal and temporal cortex of *Fmr1*^-/y^ rats (Naoki et al., 2020), and in individuals with FXS (Lovelace, Ethell, Binder, & Razak, 2018), which suggest that network-wide deficits in either gamma oscillation generation or maintenance are conserved between mammalian species.

Given the anatomical organization of inputs to the hippocampus, the power of network oscillations vary between locations within the hippocampus (Bragin et al., 1995; Buszaki, 2002; Fernández-Ruiz et al., 2017). Therefore, future work should include multisite recordings within hippocampal sub-layers and in extra-hippocampal input areas to determine the precise effects of FMRP loss on hippocampal circuit function (Radwan et al., 2016), as well as to delineate the contribution of the different brain areas that project to the hippocampus (Colgin et al., 2009).

### Weaker gamma phase locking and disrupted experience-dependent changes in gamma phase locking in CA1 pyramidal neurons of Fmr1^-/y^ rats

The spikes of pyramidal neurons in *Fmr1*^-/y^ rats were less strongly phase locked to the cycles of gamma oscillations than those of WT rats. The most robust difference in phase locking between WT and *Fmr1*^-/y^ rats was seen in the slow gamma range, consistent with observations in *Fmr1*^-/y^ mice (Talbot et al., 2018). Moreover, in WT rats, more CA1 pyramidal neurons were significantly phase locked to slow gamma on the first session in the novel environment than in subsequent sessions, but this novelty-induced increase in phase locking was not observed in *Fmr1*^-/y^ rats. Gamma oscillations in CA1 of the hippocampus reflect not only specific inputs from hippocampus (CA3) and cortex (MEC3), but also synaptic interactions between ensembles of local neurons within the hippocampus (Colgin, 2016; Csicsvari et al., 2003). As such, less phase locking to gamma in *Fmr1*^-/y^ than WT rats may not only suggest weaker coupling with extrinsic inputs, but may also reflect local microcircuit dysfunction as has been shown in the developing neocortex (Gibson et al., 2009, Domanski et al., 2019, Booker et al., 2019). Novelty-enhanced phase locking to slow gamma has been reported previously in WT rats, and requires GluA1-dependent long-term-potentiation (LTP) (Kitanishi et al., 2015). As FMRP has been proposed to promote GluA1 membrane delivery (Guo et al., 2015), and pharmacological intervention in *Fmr1*^-/y^ mice can rescue GluA1-dependent synaptic plasticity (Lim et al., 2014), it is possible that the disrupted novelty-induced gamma phase locking seen in *Fmr1*^-/y^ pyramidal neurons is due, at least in part, to abnormalities in GluA1-dependent synaptic plasticity.

### Alterations in timing of CA1 pyramidal neuron firing relative to theta oscillations

There was no difference between WT and *Fmr1*^-/y^ rats in the strength of phase locking to theta oscillations. However, there was a difference between genotypes in the phase of theta at which CA1 pyramidal neurons fired. Specifically, WT neurons fired preferentially during the late descending phase of theta in the first session in the novel environment, and their activity moved towards the early ascending phase with repeated exposures across the two days. In contrast, *Fmr1*^-/y^ pyramidal neurons fired preferentially in the early ascending phase of theta on both days and showed no experience-dependent changes.

Differences in theta phase preference between novel and familiar environments have been reported previously in WT rats (Lever et al., 2010, Fernandez-Ruiz et al., 2017, Kaefer et al., 2019). However, in those studies, novelty was associated with later firing relative to theta phase compared to a familiar environment, whereas we saw earlier theta phase firing on Day 1 than Day 2 in our novel environment in WT rats. A possible explanation for this discrepancy may relate to experimental differences, as the previous experiments compared theta phase preference between a novel and a very familiar environment; therefore, it is plausible that our environment is not sufficiently familiar on the second day of the experiment to elicit the “ familiar environment” phase preference previously reported in WT animals.

The absence of an experience-dependent change in the theta phase preference of *Fmr1*^-/y^ pyramidal neurons is consistent with a previous study in *Fmr1*^-/y^ mice (Talbot et al., 2018), and may suggest abnormal synaptic drive, arising from either excitatory or inhibitory inputs. Our *ex-vivo* data revealed that MEC3 synaptic strength is decreased in *Fmr1*^-/y^ rats. Given the known theta phase segregation of inputs to CA1 (MEC3 inputs strongest at the ascending phase and peak of the theta; Schomburg et al., 2014), based on this finding alone, we might have predicted decreased CA1 firing during the ascending phase of theta in *Fmr1*^-/y^ rats. However, the *ex-vivo* recordings also revealed that altered intrinsic excitability is able to compensate (in part) for spike output, which is consistent with our previous findings in *Fmr1*^*-/y*^ mice (Booker et al., 2020). Together, this suggests that reduced excitatory drive is compensated for within the local circuit to normalize CA1 population activity in *Fmr1*^-/y^ rats.

Given the precise temporal organization of inhibition in the hippocampus (Klausberger et al., 2003; Lapray et al., 2012) and the known abnormalities in hippocampal inhibitory transmission in the absence of FMRP (Sabanov et al., 2017), it is tempting to speculate that the absence of experience-dependent change in the theta phase preference observed in *Fmr1*^-/y^ pyramidal neurons is, at least in part, due to a persistent decrease of inhibitory tone during the ascending phase of theta in *Fmr1*^-/y^ rats, which permits pyramidal neuron discharge during that phase.

### Summary and conclusions

We report that CA1 pyramidal neurons in *Fmr1*^-/y^ rats fail to update their spatial coding of a novel environment across days of repeated experience, when compared to WT rats performing the same task. This phenomenon manifests as an inflexibility of neuronal firing properties, spatial coding, and of spike timing relative to hippocampal oscillations. Furthermore, we confirm that these changes have a plausible basis in both intrinsic cell excitability and synaptic inputs from the MEC, which may account for many of these impairments. Interestingly, absence of experience-dependent update in spatial tuning has been reported previously in a mouse model of Rett syndrome (Kee et al., 2018); while reduced spatial specificity has also been observed in two different mouse models of Down syndrome (Raveau et al., 2018; Witton et al., 2015). This raises the distinct possibility of conserved neuropathophysiology underlying cognitive abnormalities in these models, and highlights that exploring the *in-vivo* physiology of neurons in freely moving animals is critical to determining the functional consequences of neurodevelopmental disorders.

## Materials and methods

### Animals

For both *in-vivo* and *ex-vivo* studies, subjects were adult (3-4 months of age at the time of surgery) male Long-Evans Hooded WT (n = 7) and *Fmr1*^em1/PWC^ (n = 7) rats hereafter referred to as *Fmr1*^-/y^ (for more details on this rat model see [Asiminas et al., 2019]) bred in-house and kept on a 12h/12h light/dark cycle with ad libitum access to water and food prior to surgery. Following weaning (postnatal day 21) and prior to surgery, *Fmr1*^-/y^ and WT littermates were group-housed (3 to 5 rats per cage) in mixed-genotype cages. Animals were selected pseudo-randomly from a litter for use in the experiment (cohorts of 2-6 animals at a time). Each cohort included both WT and *Fmr1*^-/y^ rats. This was done by an experimenter not involved in surgery, data collection or analysis, by randomly picking rat ID numbers from a given litter (while ensuring balance of WT and KO rats). Experimenters involved in data collection and data analysis were blind to the genotype of the subjects throughout all stages of the experiments and spike sorting until final statistical analyses were conducted [in line with the ARRIVE (Animal Research: Reporting of *In-Vivo* Experiments) guidelines (du Sert et al., 2020). Following implantation surgeries, rats were housed individually in cages designed to minimize head-stage damages after surgery.

All recordings were performed during the light phase of the cycle. Following surgery and recovery, rats were food restricted such that they maintained approximately 90% of their free-feeding weight. For *ex-vivo* slice electrophysiology, a total of 28 rats (14 WT, 14 *Fmr1*^*-/y*^) were used at 2-3 months of age. This experiment complied with the national [Animals (Scientific Procedures) Act, 1986, United Kingdom] and international [European Communities Council Directive of November 24, 1986 (86/609/EEC)] legislation governing the maintenance of laboratory animals and their use in scientific experiments. Local ethical approval was granted by the University of Edinburgh Animal Welfare and Ethical Review Board.

### Electrodes and surgery

The microdrives used in this study were based on a modified tripod design described previously (Kubie, 1984). The drives were loaded with eight tetrodes, each of which was composed of four HML coated, 17 µm, 90% platinum 10% iridium wires (California Fine Wire, Grover Beach, CA). Tetrodes were threaded through a thin-walled stainless steel cannula (23 Gauge Hypodermic Tube, Small Parts Inc, Miramar, FL). The tip of every wire was gold-plated (Non-Cyanide Gold Plating Solution, Neuralynx, MT) to reduce the impedance of the electrode from a resting impedance of 0.7–0.9 MΩ to a plated impedance in the range of 150–250 kΩ (200 kΩ being the target impedance) one to ten hours before surgery. Electrodes were implanted using standard stereotaxic procedures under isoflurane anaesthesia. Hydration was maintained by subcutaneous administration of 2.5 ml 5% glucose and 1 mL 0.9% saline. Animals were also given an anti-inflammatory analgesia (small animal Carprofen/Rimadyl, Pfizer Ltd., UK) subcutaneously. Electrodes were lowered to just above the dorsal CA1 cell layer of the hippocampus (−3.5 mm AP from bregma, +2.4 mm ML from the midline, −1.7 mm DV from dura surface). The drive assembly was anchored to the skull screws and bone surface using dental acrylic (Associated Dental Products Ltd. Swindon, UK). Animals were monitored closely for at least two hours in their home cage while recovering from anaesthesia, and then returned to the colony. Following this, at least one week of recovery time passed before access to food was restricted and screening for cellular activity began.

### Unit recording

Single unit and local field potential (LFP) activity were recorded using a 32-channel Axona USB system (Axona Ltd., St. Albans, UK). Mill-Max connectors built into the rat’s microdrive were attached to the recording system via two unity gain buffer amplifiers and a light, flexible, elasticated recording cable. The recording cable passed signals through a ceiling mounted slip-ring commutator (Dragonfly Research and Development Inc., Ridgeley, West Virginia) to a pre-amplifier where they were amplified 1000 times. The signal was then passed to a system unit; for single unit recording the signal was band-pass (Butterworth) filtered between 300 and 7000 Hz. Signals were digitized at 48 kHz (50 samples per spike, 8 bits/sample) and could be further amplified 10–40 times at the experimenter’s discretion. The LFP signals were recorded from one channel of a tetrode located in the pyramidal neuron layer of the dCA1 area. The signals were amplified by a factor of 1000–2000, low-pass filtered at 500 Hz, and sampled at a rate of 4.8 kHz (16 bits/sample). A notch filter was applied at 50 Hz. The position of the animal was recorded by tracking two small light-emitting diodes fixed on the headstage connected to the rat’s microdrive. A ceiling mounted, infrared sensitive CCTV camera tracked the animal’s position at a sampling rate of 50 Hz. Rats were screened for single unit activity and for the presence of theta oscillations once or twice a day, at least five days a week, while foraging for chocolate treats (CocoPops, Kellogg’s, Warrington, UK) in a blue, wooden square recording environment (1 m × 1 m × 52 cm) which was not used during later experiments. At the end of each screening session, rats were removed from the recording apparatus and the electrodes were lowered if no hippocampal unit activity had been observed. Brain tissue was allowed to recover from electrode movement for at least 5 hours before a new screening session started.

### Recording protocol

Once at least 10 putative pyramidal neurons were detected, rats were transferred to a totally novel grey plastic cylindrical environment (62 cm diameter, 60 cm walls) surrounded by black curtains within the same experimental room. The cylinder contained one salient black cue card that remained stable throughout the two days of recording. Three 10-min recording sessions, separated by a 10 min inter-session interval (ISI), took place on each of the two days of the experiment, during which rats foraged for scattered chocolate treats in the recording arena (six sessions in total) (Figure 1). Between sessions of the same day, rats were placed in a plastic holding bucket (25 cm diameter, sawdust bedding) while remaining tethered.

### Single Unit analysis

Single unit activity was analysed offline using a custom-written MATLAB (MathWorks) routine that makes use of the Klustakwik spike sorting program (Harris, Henze, Csicsvari, Hirase, & Buzsáki, 2000). The dimensionality of the waveform information was reduced to the first principal component, energy, peak amplitude, peak time, and width of the waveform. The energy of a signal *x* was defined as the sum of squared moduli given by the formula:

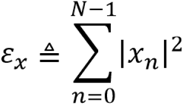

Based on these parameters, Klustakwik spike sorting algorithms were then used to distinguish and isolate separate clusters. The clusters were then further checked and refined manually using the manual cluster cutting GUI, Klusters (Hazan, Zugaro, & Buzsáki, 2006). As well as the previously mentioned features, manual cluster cutting also made use of spike auto- and cross-correlograms to examine refractory period, complex spiking and potential theta modulation. Cluster quality was operationalized by calculating isolation distance (Iso-D), L_ratio_, and peak waveform amplitude, taken as the highest amplitude reached by the four mean cluster waveforms. For cluster *C*, containing *n*_*c*_ spikes, Iso-D is defined as the squared Mahalanobis distance of the *n*_*c*_*-th* closest *non-c* spike to the centre of *C*. The squared Mahalanobis distance was calculated as:

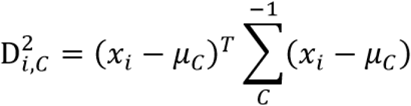

where *x*_*i*_ is the vector containing features for spike *i*, and *μ*_*C*_ is the mean feature vector for cluster *C*. A higher value indicates better isolation from non-cluster spikes (Schmitzer-Torbert, Jackson, Henze, Harris, & Redish, 2005). The L quantity was defined as:

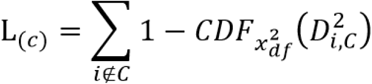

where i∉C is the set of spikes which are not members of the cluster and *CDF* is the cumulative distribution function of the distribution with 8 degrees of freedom. The cluster quality measure, *L*_*ratio*_ was thus defined as *L* divided by the total number of spikes in the cluster (Schmitzer-Torbert & Redish, 2004).

A cluster was classified as a pyramidal neuron on the maze if it satisfied the following criteria: i) Iso-D >15 and L_ratio_ <0.2, ii) the width of the waveform was >250 μs, and iii) the mean firing rate on the maze was greater than 0.1 Hz but less than 5Hz. For a pyramidal neuron to be classified as place cells, its spatial information content should be greater than 0.5 bit/spike. Spatial information content is given by the equation:

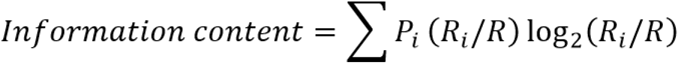

where *i* is the bin number, *Pi* is the probability for occupancy of bin *i, Ri* is the mean firing rate for bin *i*, and *R* is the overall average firing rate (Skaggs, McNaughton, Gothard, & Markus, Etan, 1993).

To assess the stability of spatial discharge patterns of spatially modulated pyramidal neurons, we calculated the Pearson correlation between firing rate maps of successive recording sessions. Firing rate maps were produced for each session by dividing the recording cylinder area into a grid of 2.5 cm square bins. The firing rate in each bin was calculated as the total number of spikes which occurred in that bin divided by the total length of time spent there. Bins in which the rats spent less than 100ms were treated as if they had not been visited. Bins were smoothed using a Gaussian filter with the following parameters:

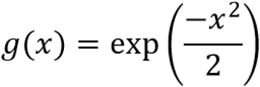

These bin-specific firing rates were plotted in a heat map, showing where the preferred firing location of a cell was in a given environment.

The algorithm for calculating firing rate is then given by:

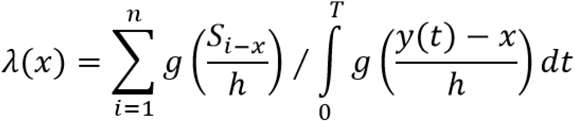

where *S*_*i*_ represents the positions of every recorded spike, *x* is the centre of the current bin, the period [0 T] is the recording session time period, *y(t)* is the position of the rat at time *t*, and *h* is a smoothing factor, which was set to 2.5 cm.

### Spectral Analysis

Position data from each session were binned into 500ms epochs and the velocity for each epoch was calculated. Raw LFP traces (4.8 KHz) were z-scored (mean was subtracted and divided by standard deviation). LFP data from periods of immobility (<3 cm/s) were excluded from subsequent analysis. Time-frequency spectrograms were calculated using Chronux Toolbox (http://chronux.org; Bokil, Andrews, Kulkarni, Mehta, & Mitra, 2010), function *mtspecgramc()* using a window size and time step of 20 s and 10 s, respectively (Schlesiger et al., 2015). Power estimates for the frequency bands of interest [Theta (6-12 Hz), Slow Gamma (30-50 Hz), Medium Gamma (60-100 Hz)] were excised from the spectrogram and averaged. The average value over all the 4 s periods of activity (>3 cm/s) following periods of immobility is represented for all frequencies in Fig. 6C. For the relationships between running speed and oscillation power, EEG signals from all 500 ms epochs were stratified based on the velocity (3 cm/s wide bins). Power spectra estimation during each epoch was done by means of the Welch periodogram method (50% overlapping Hamming windows), which was obtained by using the *pwelch()* function from MATLAB Signal Processing Toolbox. Specific band powers were computed by integrating the power spectral density (PSD) estimate for each frequency range of interest [MATLAB function *bandpower()*].

### Phase-locking analysis

To investigate spike timing along oscillations, a band-pass filter was applied to the LFP signals. The low cut-off stop band was the low passband minus 2 Hz; the high cut-off stop band was the high passband plus 2 Hz [Theta (4-14 Hz), Slow Gamma (28-52 Hz), Medium Gamma (58-102 Hz)]. Both the instantaneous amplitude and the phase time series of a filtered signal were computed from the Hilbert transform, which was obtained by using the *hilbert()* function from the MATLAB Signal Processing Toolbox. Only spikes during windows (400 ms) of strong oscillations (>2 standard deviations of mean power) were included in this analysis (Colgin et al., 2009; Kitanishi et al., 2015). Every recorded spike from these periods was assigned a spike phase *θ*_*j*_, where *j* denotes the *j-th* spike. The mean resultant vector r was calculated as:

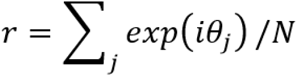

where *N* is the total number of spikes. The strength of phase locking (resultant length) was defined as |r|. Theoretically, this value ranges from 0 to 1. The value is zero if the phases are uniformly distributed along the phases of gamma oscillations, while it is one if all spikes fire at the exactly same phase. In practice, the values for individual neurons are distributed mostly in the range of 0-0.2 as shown in Figure 7. The trough of gamma oscillation was defined as 0/360°.

### Histology

At the end of the experiment animals were given an overdose of pentobarbital intraperitoneally (Euthatal, Merial Animal Health Ltd., Essex, UK), and perfused with 0.9% saline solution followed by a 4% formalin solution. The brain was extracted and stored in 4% formalin for at least seven days prior to any histological analyses. The brains were sliced in 32 µm sections on a freezing microtome at −20°. These sections were stained with a 0.1% Cresyl violet solution and the tissue section that best revealed the electrode track was imaged using ImageJ software (ImageJ, NIH, Bethesda).

### *Ex-vivo* electrophysiology

*Ex-vivo* brain slices were prepared as previously described (Booker, 2020). Briefly, rats were sedated with isoflurane, anaesthetized with sodium pentobarbital (100 mg/kg) and then transcardially perfused with ice-cold, carbogenated (95 % O2/5 % CO2), and filtered sucrose-modified artificial cerebrospinal fluid (sucrose-ACSF; in mM: 87 NaCl, 2.5 KCl, 25 NaHCO3, 1.25 NaH2PO4, 25 glucose, 75 sucrose, 7 MgCl2, 0.5 CaCl2). Once perfused, the brain was rapidly removed and 400 μm slices containing the dorsal pole of the hippocampus were cut on an oscillating blade vibratome (VT1200S, Leica, Germany) in the coronal and plane. Slices were then transferred to a submerged chamber containing sucrose-ACSF at 35 °C for 30 min, then at room temperature until needed.

For recording, slices were transferred to a submerged chamber perfused with pre-warmed carbogenated ACSF (in mM: 125 NaCl, 2.5 KCl, 25 NaHCO3, 1.25 NaH2PO4, 25 glucose, 1 MgCl2, 2 CaCl2) at a flow rate of 4-6 mL.min-1 at 31± 1 °C) which contained 50 µM picrotoxin to block GABA_A_ receptor mediated currents. Neurons were visualized under infrared differential inference contrast (IR-DIC) microscopy with a digital camera (SciCamPro, Scientifica, UK) mounted on an upright microscope (SliceScope, Scientifica, UK) with a 40x water-immersion objective lens (1.0 N.A., Olympus, Japan). Whole-cell patch-clamp recordings were performed with a Multiclamp 700B (Molecular Devices, CA, USA) amplifier, using recording pipettes pulled from borosilicate glass capillaries (1.7 mm outer/1 mm inner diameter, Harvard Apparatus, UK) on a horizontal electrode puller (P-97, Sutter Instruments, CA, USA). Pipettes were filled with a K-gluconate based internal solution (in mM 142 K-gluconate, 4 KCl, 0.5 EGTA, 10 HEPES, 2 MgCl_2_, 2 Na_2_ATP, 0.3 Na_2_GTP, 1 Na_2_Phosphocreatine, 2.7 Biocytin, pH=7.4, 290-310 mOsm) which gave a 3-5 MΩ tip resistance. Neurons were rejected if: they were more depolarized than -50 mV, had an access resistance >30 MΩ, or the access resistance changed by more than 20% during the recording. Cell-attached recordings were performed as above, but without breaking through into the whole-cell configuration.

Intrinsic membrane properties were measured in current-clamp. Passive membrane properties, including resting membrane potential, membrane time constant, input resistance, and capacitance were measured from small hyperpolarizing current steps (10 pA, 500 ms duration), from a zero-current level. Active properties were determined from a series of depolarizing current steps (0 to +400 pA, 500 ms) from -70 mV, maintained by addition of bias current. AP properties were determined from the first, second or fifth AP elicited at rheobase. Stimulation of the Schaffer-Collateral (SC) and Temporoammonic (TA) pathways were made with a bipolar twisted Ni:Chrome wire electrode placed in either *str. radiatum* or *str. lacunosum-moleculare* in distal CA1. In all stimulation slices, CA3 was severed to prevent recurrent activation from antidromic activation of CA3. To assess synaptic strength of afferent inputs, 2x stimuli of 200 µs duration (50 ms interval) were delivered at 10 second intervals at 30 V, 60 V, and 90 V levels from a constant-voltage stimulation box (Digitimer, Cambridge, UK). Recordings were first performed in cell-attached mode to identify cell spike output. Following breakthrough into whole-cell mode, EPSPs were recorded in current-clamp configuration with membrane potential biased to -70 mV. All recordings were filtered online at 10 kHz with the built-in 4-pole Bessel Filter and digitized at 20 kHz (Digidata1440, Molecular Devices, CA, USA). Traces were recorded in pCLAMP 9 (Molecular Devices, CA, USA) and stored on a personal computer. Analysis of electrophysiological data was performed offline using the open source software package Stimfit (Guzman, Schlögl, and Schmidt-Hieber 2014), blind to both genotype and treatment conditions.

### Axon initial segment (AIS) labelling and neuron visualization

Additional *ex-vivo* brain slices were collected during preparation of tissue for *ex-vivo* recordings (see above) and fixed for 1 hour at room temperature in 4% paraformaldehyde in 0.1 M phosphate buffer (PB). Following fixation, slices were transferred to 0.1 M phosphate buffered saline (PBS) and stored for up to 1 week. Immunohistochemistry was performed as previously described (Oliveira et al., 2021). Briefly, slices were rinsed 3-4 times in PBS, then blocked in a solution containing 10% normal goat serum, 0.3% Triton X-100 and 0.05% NaN_3_ diluted in PBS for 1 hour. Primary antibodies raised against AnkyrinG (1:1000; 75-146, NeuroMab, USA) and NeuN (1:1000, Millipore EMD, UK) were applied in a solution containing 5% normal goat serum, 0.3% Triton X-100 and 0.05% NaN_3_ diluted in PBS, for 24-72 hours at 4 °C. Slices were then washed in PBS and secondary antibodies (AlexaFluor 488 and AlexaFluor 633, Invitrogen, UK, both 1:500) were applied in a solution containing 3% normal goat serum, 0.1% Triton X-100 and 0.05% NaN_3_ overnight at .4°C. Slices were rinsed in PBS, desalted in PB and mounted on glass slides with Vectashield Hard-Set mounting medium (Vector Labs, UK). Stacks of images of the lower *str. pyramidale* upper *str. oriens* were acquired on a Zeiss LSM800 laser scanning confocal microscope, under a 60x (1.2 NA) objective lens at 2048×2048 resolution, with a step size of 0.25 µm. AIS lengths were measured offline using ImageJ as segmented lines covering the full extent of AnkyrinG labelling observed. A minimum of 25 AIS were measured for each rat.

For CA1 pyramidal neuron reconstructions, fixed slices containing recorded neurons were fixed overnight in 4% paraformaldehyde + 0.1 M PB at 4 °C. Slices were then transferred to 0.1 M PB and stored until processing. For visualization, slices were washed 2-3 times in 0.1 M PB then transferred to a solution containing Streptavidin conjugated to AlexaFluor568 (1:500, Invitrogen, UK) vand 0.3% Triton X-100 and 0.05% NaN_3._ Slices were then incubated for 48-72 hours at 4 °C. Slices were then washed in 0.1 M PB and mounted on glass slides with an aqueous mounting medium (VectaShield, Vector Labs, UK) and coverslipped. Neurons were imaged on an upright confocal (as above) with image stacks collected with a 20x objective lens (2048×2048, 1 µm steps). Neurons were reconstructed with the SNT toolbox for FIJI/ImageJ (Arshadi et al., 2020), Sholl analysis performed, and branch lengths measured for the different dendritic compartments.

Dendritic protrusion analysis was performed on short dendritic segments (secondary dendrites) that were imaged with a 63x objective lens (2.4x digital zoom, 2048×2048 resolution giving 40 nm pixels, 0.13 µm z-step). These images were deconvolved (Huygens Software Package, Scientific Volume Imaging, The Netherlands), then dendritic protrusions counted as a function of length in FIJI. To estimate the total number of dendritic protrusions per dendritic compartment, the density was multiplied by the total length of dendrites in that compartment.

### Statistical Analyses

#### In-vivo electrophysiology

We compared the firing properties of the identified pyramidal neurons across days and sessions in WT and *Fmr1*^-/y^ rats in three ways: first, we modelled our data using a generalized linear mixed model (glmm) approach in order to take into account the hierarchy of dependency in our data sets (genotypes-rats-neurons) and account for random effects; second, we analysed the distribution of values for each property examined across all neurons; finally we analysed the neuronal properties at the rat level by calculating the average value for each property across the neurons recorded in each rat.(for additional discussion of the choice of statistical approach, see Leger and Didrichsons, 1994, Editorial JNeurosci 2018).

Statistical modelling routines for the Linear Mixed-Effects (LME) models were written and run using RStudio 1.0.153 (RStudio Team, 2016). Depending on the data distribution, Linear Mixed Models (LMMs) or Generalized Linear Mixed Models (GLMMs) were fitted to single unit data metrics using the R package lme4 v1.1-17 (Bates, Mächler, Bolker, & Walker, 2015). Animal and cell identity was included in models as a random effect, and the variables (terms) of interest (genotype, day, session-in-day) along with all interactions between them were included as fixed effects. Interactions and terms are progressively eliminated when a simpler model (a model not containing that term or interaction) fits the data equally well (based on likelihood ratio test). Consequently, the p-values reported in the context of LMEs are given by likelihood ratio tests between a model containing the variable or interaction in question and a model without that variable or interaction (a reduced/null model).

For rat average data analysis as well as for band oscillatory power changes, we used a three-way ANOVA statistical analysis (genotype x day x session); the normality of the data was assumed and limitations of this are acknowledged. For analysis of the velocity effects on oscillatory power (Figure 6 Supp 1) we used a three-way ANOVA statistical analysis (genotype x day x velocity).

The percentage of phase locked cells by each oscillatory band (Figure 7 H-J) were analysed by fitting a series of multiple logistic regression models. For each dataset, a multiple logistic regression model was fit to the data with genotype, session and their interaction: Phaselock ∼ Intercept + Genotype + Session + Genotype: Session (Model 1). The fit of Model 1 was compared using Akaike’s Information Criterion to the fit of a simpler model that did not contain the interaction term: Phaselock ∼ Intercept + Genotype + Session (Model 2). In the case that Model 1 (model containing interaction) was the preferred model, the two-proportion z-test was used to explore the data (see below). To explore main effects of genotype and session, two separate models were fitted: Phaselock ∼ Intercept + Genotype (Model 3) and Phaselock ∼ Intercept + Session (Model 4) and the null hypothesis that the simplest (intercept-only) model is correct was tested again using Log-likelihood ratio (G squared). For individual comparisons between genotypes and sessions, we used two-proportion z-test (Kitanishi et al., 2005) with correction for false discovery rate (alpha = 0.05) with Benjamini-Hochberg procedure.

As the glmm framework for analysis of circular data is still under development, for analysis of oscillatory phase preference (Figure 8), we first used Harrison-Kanji test (Harrison & Kanji, 1988) to analyse effects of genotype, day as well as the interaction between them with neuron as the unit of measurement. For comparisons between groups and days, we used the Watson-Williams test (Watson & Williams, 1956).

Statistical evaluation of data is presented in figure legends, main manuscript and supplementary tables. Statistical comparisons between distributions of correlations (r) were performed on Fisher z-transformed correlation values: z=(0.5) 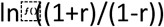. Average ± SEM values are reported throughout the manuscript unless stated otherwise. Significance was set at p<0.05. Statistical analysis was carried out in SPSS 16.0 (IBM), R v.3.4.4 (R Core Team, 2018) or MATLAB (CircStat MATLAB toolbox) (Berens 2009), and graphs created in GraphPad (Prism 8).

#### Ex-vivo electrophysiology and anatomical analysis

Data generated from *ex-vivo* brain slices was analysed as described previously (Booker *et al*., 2020). Group sizes were chosen based on a presumed effect size of 15% and an overall statical power of 80% (N=7-8/group). For assessment of genotype effect, data were analysed using an LME based approach (see above), with animal and slice identity set as random effects. The p-value was then approximated using the Wald test, with effect size and variation estimated from the LMEs fitted to the data. For synaptic stimulation experiments, either the amplitude of the EPSP or the spike-probability were plotted per slice and compared using a 2-way ANOVA. If a genotype/stimulus interaction was observed, then Sidak *posthoc* tests were performed (corrected for multiple comparisons). For AIS lengths and morphology analysis, an LME analysis was performed, again using animal and slice identity as random effects with p-value approximated using the Wald test. For dendritic protrusion analysis, the animal average was analysed with Student’s t-test

## Figure Supplements

**Figure 1 - Supplement 1.**
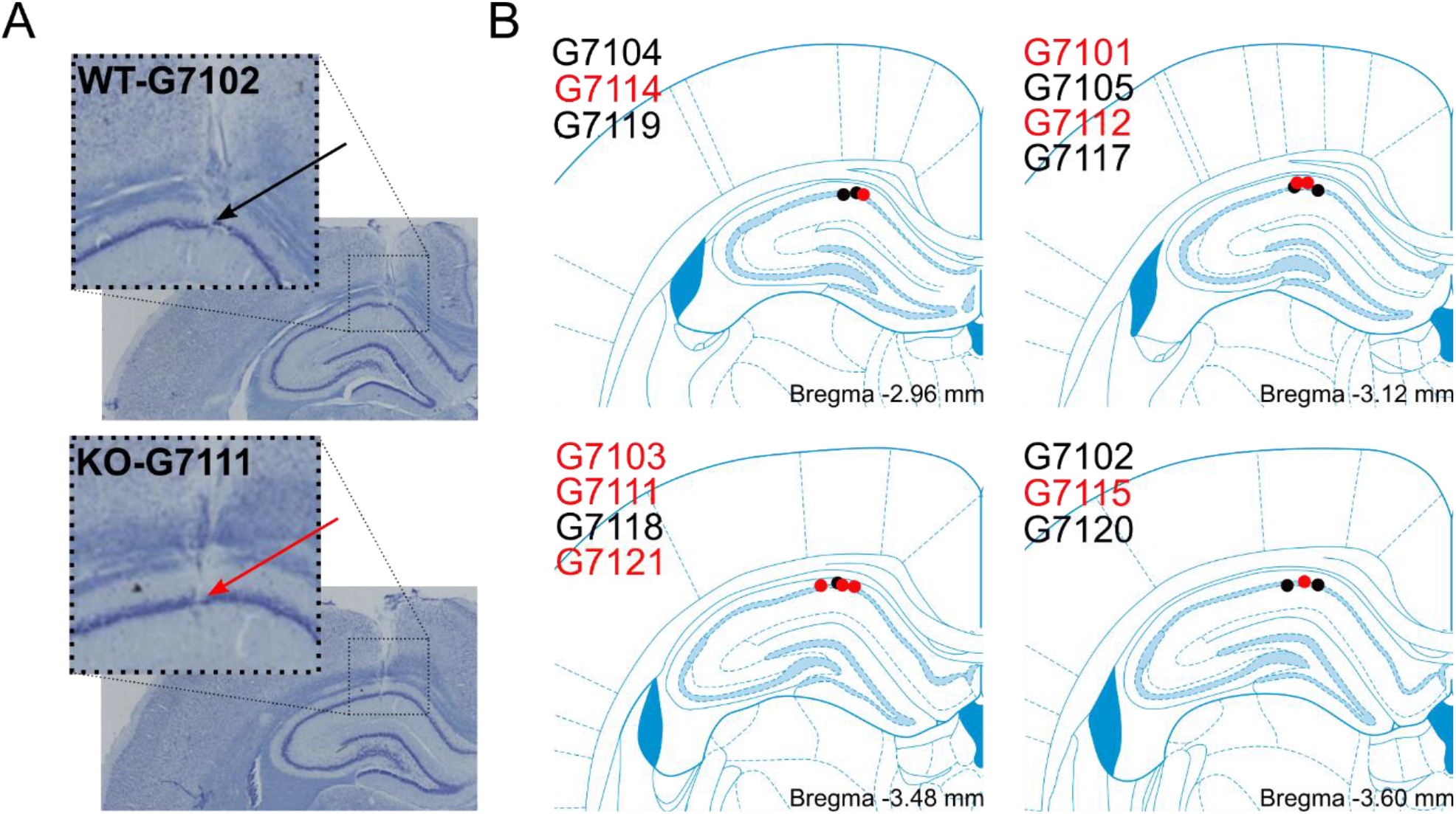
Histological confirmation of electrode placement. **(A)** Example coronal brain sections from a WT and an *Fmr1*^*-/y*^ rat stained with cresyl violet. Arrows indicate the recording locations from the pyramidal neuron layer in the WT (black) and *Fmr1*^*-/y*^ (red) rat. **(B)** Schematics of individual electrode placements in the CA1 cell layer of the hippocampus from all rats. Dots represent the location of the tip of the tetrode bundle at the end of recording. Each dot is labelled with a rat number and dot colour indicates genotype (black for WT and red for *Fmr1*^*-/y*^). The four different schematics reflect the estimated anterior-posterior (AP) coordinate (AP -3.48 mm from bregma being the intended coordinate).

**Figure 1 - Supplement 2.**
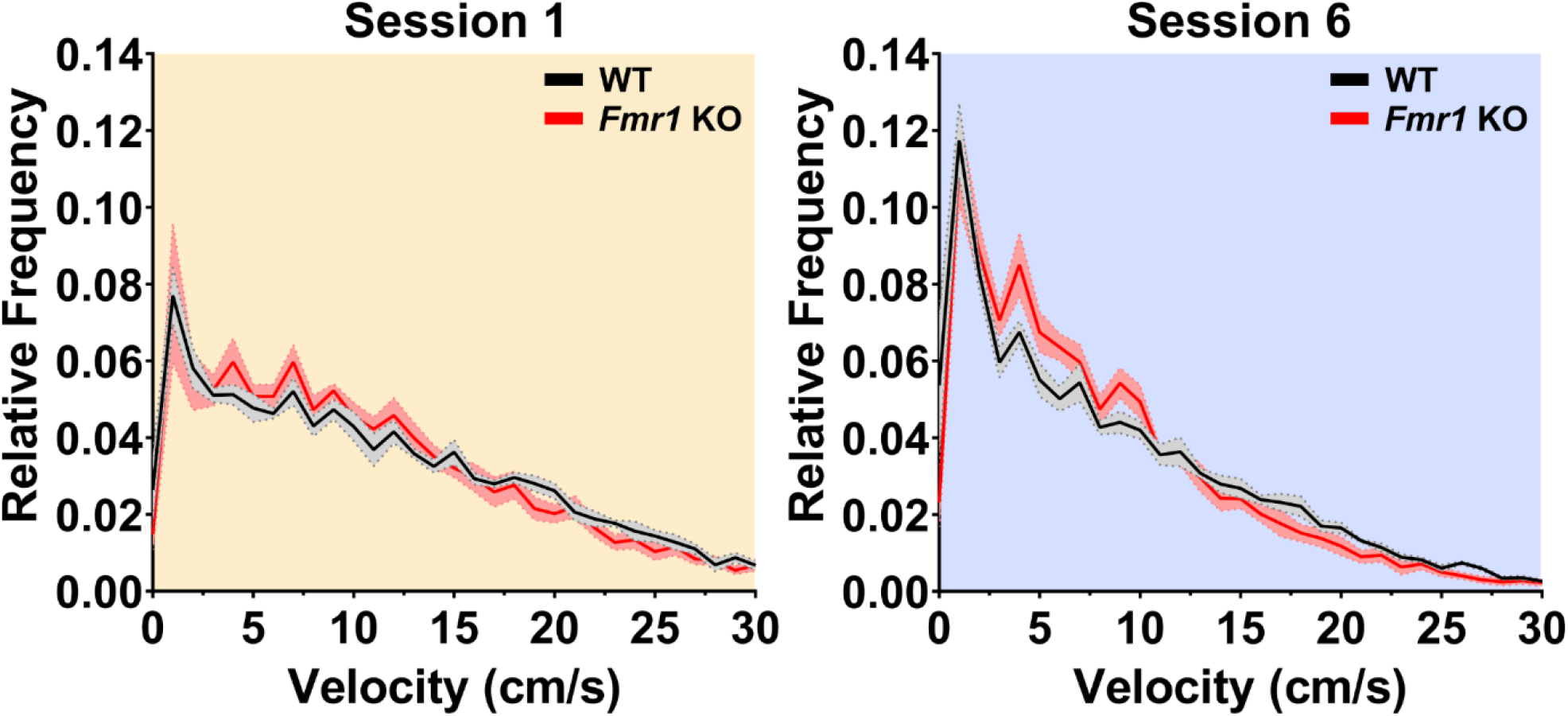
Ambulatory activity decreases over repeated exploration of an initially novel environment for both WT and *Fmr1*^-/y^ rats. Velocity distributions for 500 ms epochs in the first and last recording session for WT and *Fmr1*^-/y^ rats did not reveal any differences between groups. Solid lines depict rat means; shaded areas depict SEM. Pale yellow and pale purple backgrounds denote data from Day 1 and Day 2 respectively.

**Figure 1 - Supplement 3.**
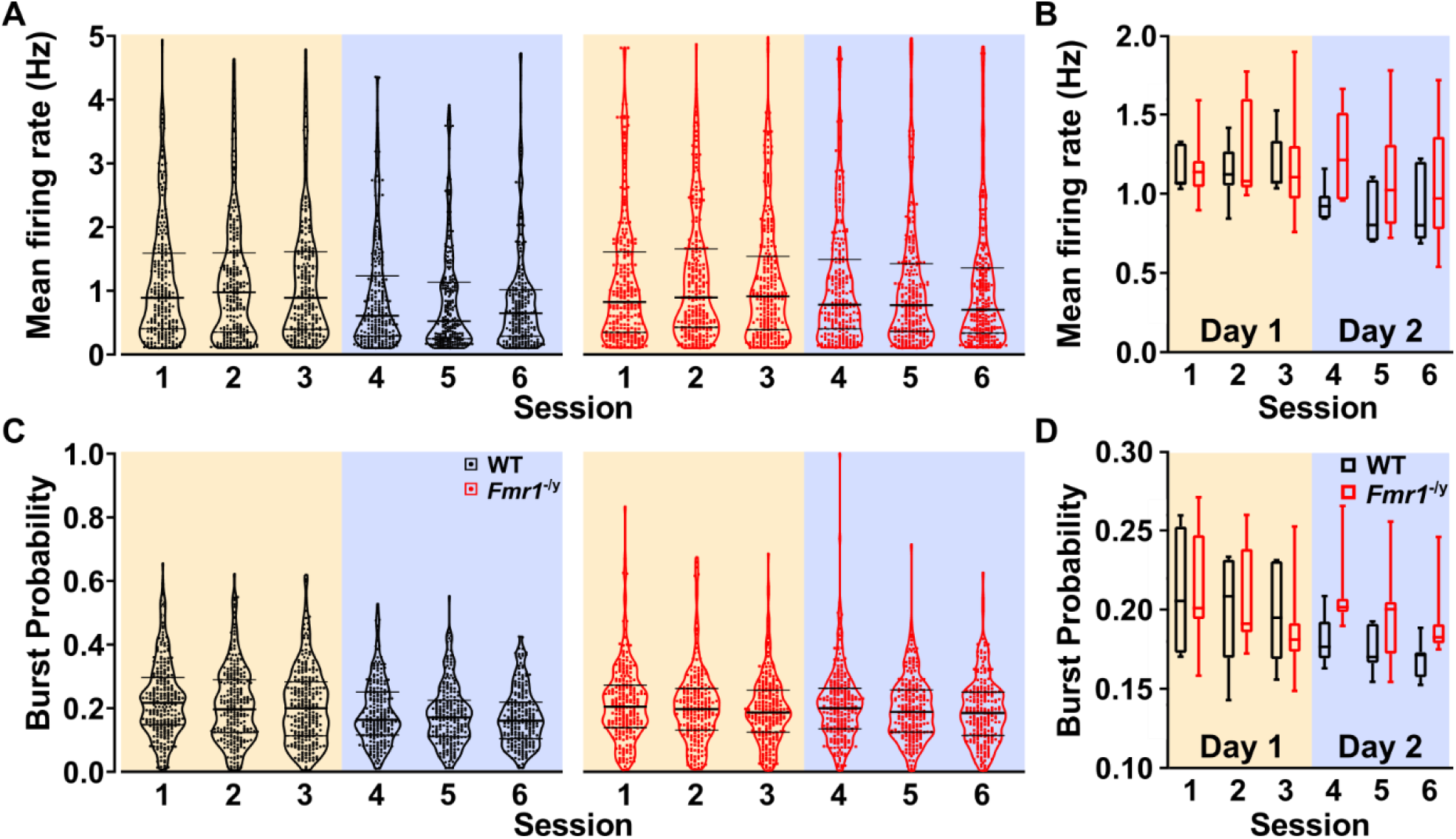
**(A)** Violin plots depicting the mean firing rate distributions across all six recording sessions for WT and *Fmr1*^*-/y*^ pyramidal neurons. **(B)** Mean firing rate plotted and analysed at the rat average level. Three-way RM ANOVA: Day x Session x genotype: F_(2,24)_=0.788, p=0.466; Day x Genotype: F_(1,12)_=1.949, p=0.188; Session x Genotype: F_(2,24)_=1.089, p=0.353; Genotype: F_(1,12)_=1.147, p=0.305; Day: F_(1,12)_=5.933, p=0.031; Session: F_(2,24)_=0.674, p=0.532. **(C-D)** Same as (**A**-**B**) for burst probability. Three-way RM ANOVA: Day x Session x genotype: F_(2,24)_=0.427, p=0.657; Day x Genotype: F_(1,12)_=7.005, p=0.021; Session x Genotype: F_(2,24)_=0.246, p=0.784; Genotype: F_(1,12)_=1.060, p=0.324; Day: F_(1,12)_=6.165, p=0.029; Session: F_(2,24)_=6.109, p=0.007. *Posthoc* tests Day1: WT vs *Fmr1*^*-/y*^ p=0.974; Day2: WT vs *Fmr1*^*-/y*^ p=0.041; WT Day1 vs Day2 p=0.034; *Fmr1*^*-/y*^ Day1 vs Day2 p=0.820. For box and violin plots the middle line represents rat median, upper and lower end of the boxes, and upper and lower line in the violins represent 95^th^ and 5^th^ percentile, error bars in box plots represent maximum and minimum values. N_WT_ = 7, N_KO_ =7. Pale yellow and pale purple backgrounds denote data from Day 1 and Day 2 respectively.

**Figure 1 – Supplement Table 1.**
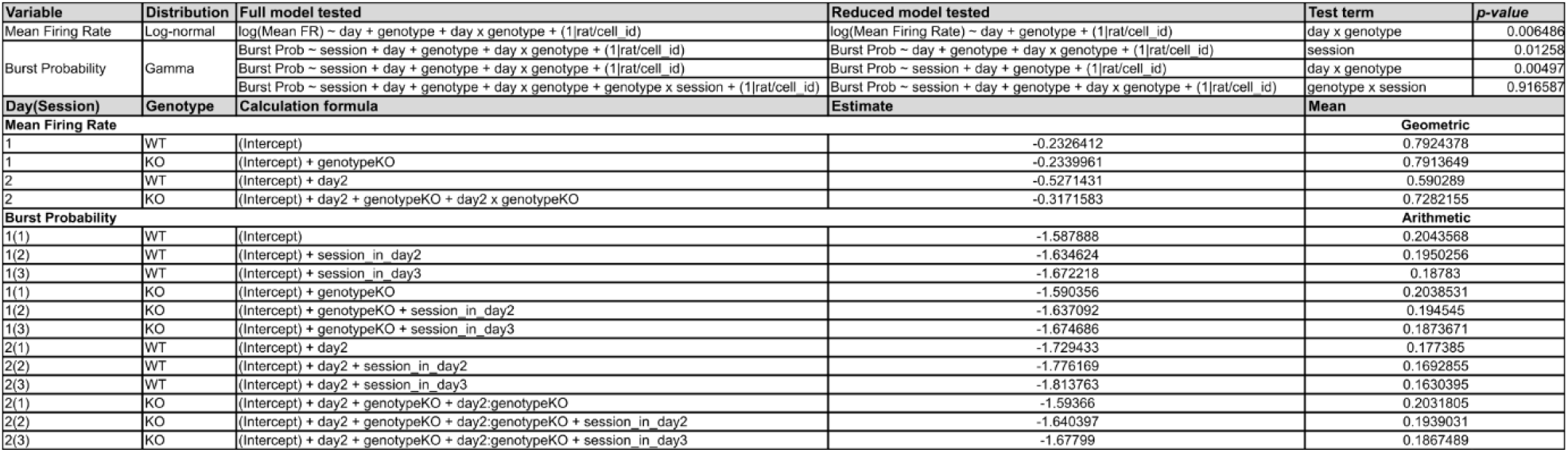
LME modelling estimates

**Figure 2 - Supplement 1.**
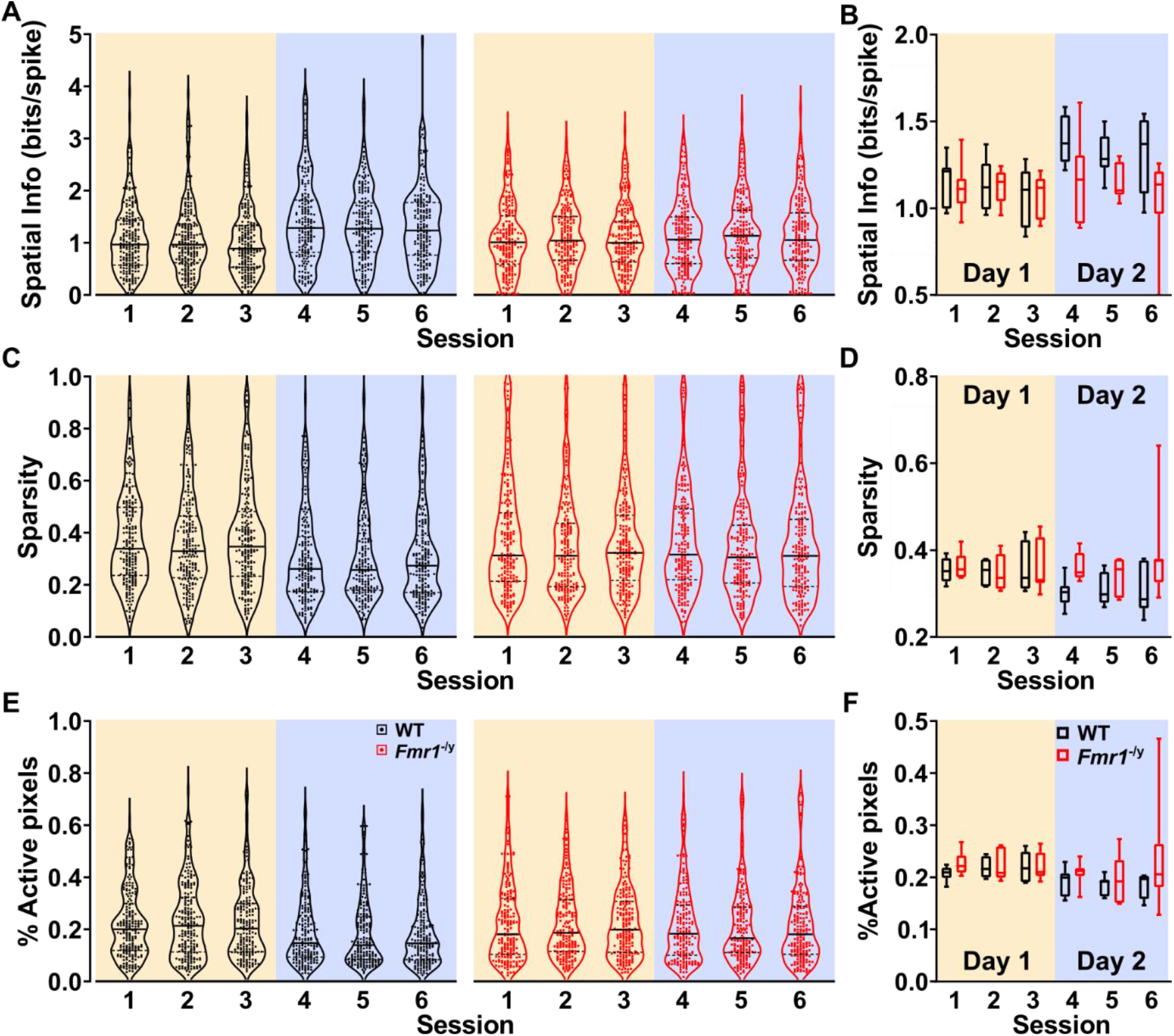
**(A)** Violin plots depicting the spatial information distributions across all six recording sessions for WT and *Fmr1*^*-/y*^ pyramidal neurons. **(B)** Spatial information plotted and analysed at the rat average level. Three-way RM ANOVA: Day x Session x genotype: F_(2,24)_=0.561, p=0.578; Day x Genotype: F_(1,12)_=7.363, p=0.019; Session x Genotype: F_(2,24)_=0.211, p=0.811; Genotype: F_(1,12)_=2.875, p=0.116; Day: F_(1,12)_=9.285, p=0.010; Session: F_(2,24)_=2.132, p=0.141. *Posthoc* tests Day1: WT vs *Fmr1*^*-/y*^ p=0.992; Day2: WT vs *Fmr1*^*-/y*^ p=0.023; WT Day1 vs Day2 p=0.002; *Fmr1*^*-/y*^ Day1 vs Day2 p=0.846. **(C-D)** Same as (**A**-**B**) for sparsity. Three-way RM ANOVA: Day x Session x genotype: F_(2,24)_=0.762, p=0.478; Day x Genotype: F_(1,12)_=8.472, p=0.013; Session x Genotype: F_(2,24)_=0.687, p=0.513; Genotype: F_(1,12)_=2.460, p=0.143; Day: F_(1,12)_=5.189, p=0.042; Session: F_(2,24)_=1.364, p=0.275. *Posthoc* tests Day1: WT vs *Fmr1*^*-/y*^ p=0.948; Day2: WT vs *Fmr1*^*-/y*^ p=0.018; WT Day1 vs Day2 p<0.001; *Fmr1*^*-/y*^ Day1 vs Day2 p=0.745. **(E-F)** Same as (**A**-**B**) for %Active pixels. Three-way RM ANOVA: Day x Session x genotype: F_(2,24)_=0.719, p=0.497; Day x Genotype: F_(1,12)_=2.103, p=0.173; Session x Genotype: F_(2,24)_=0.691, p=0.511; Genotype: F_(1,12)_=2.560, p=0.136; Day: F_(1,12)_=6.977, p=0.022; Session: F_(2,24)_=0.687, p=0.513. For box and violin plots the middle line represents rat median, upper and lower end of the boxes, and upper and lower line in the violins represent 95^th^ and 5^th^ percentile, error bars in box plots represent maximum and minimum values. N_WT_ = 7, N_KO_ =7. Pale yellow and pale purple backgrounds denote data from Day 1 and Day 2 respectively.

**Figure 2 – Supplement Table 1.**
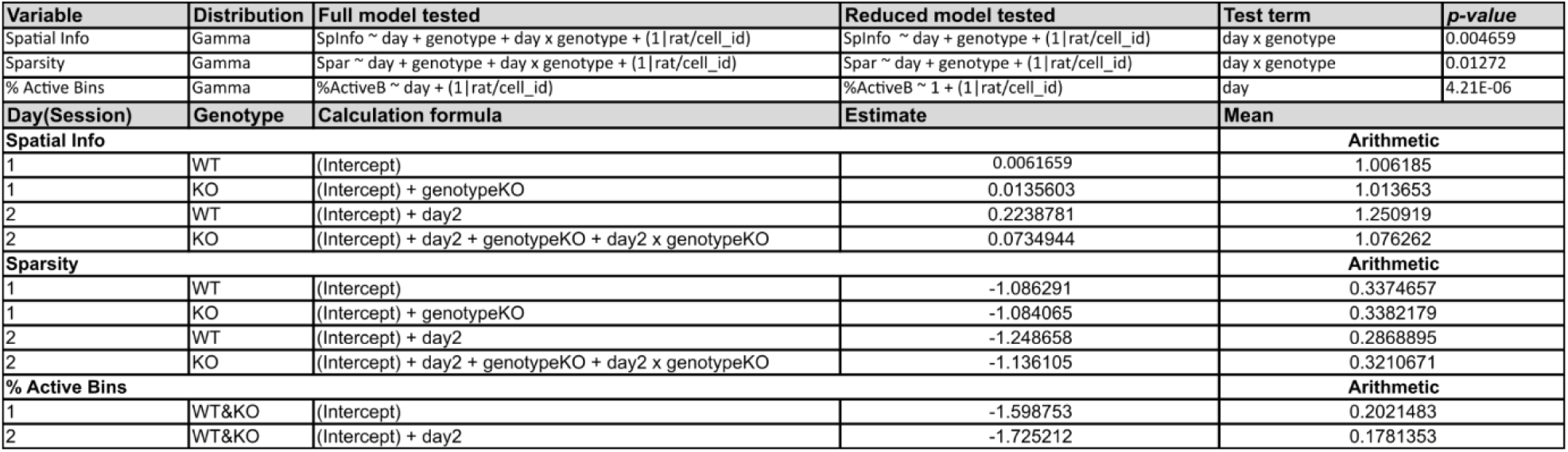
LME modelling estimates

**Figure 3 - Supplement 1.**
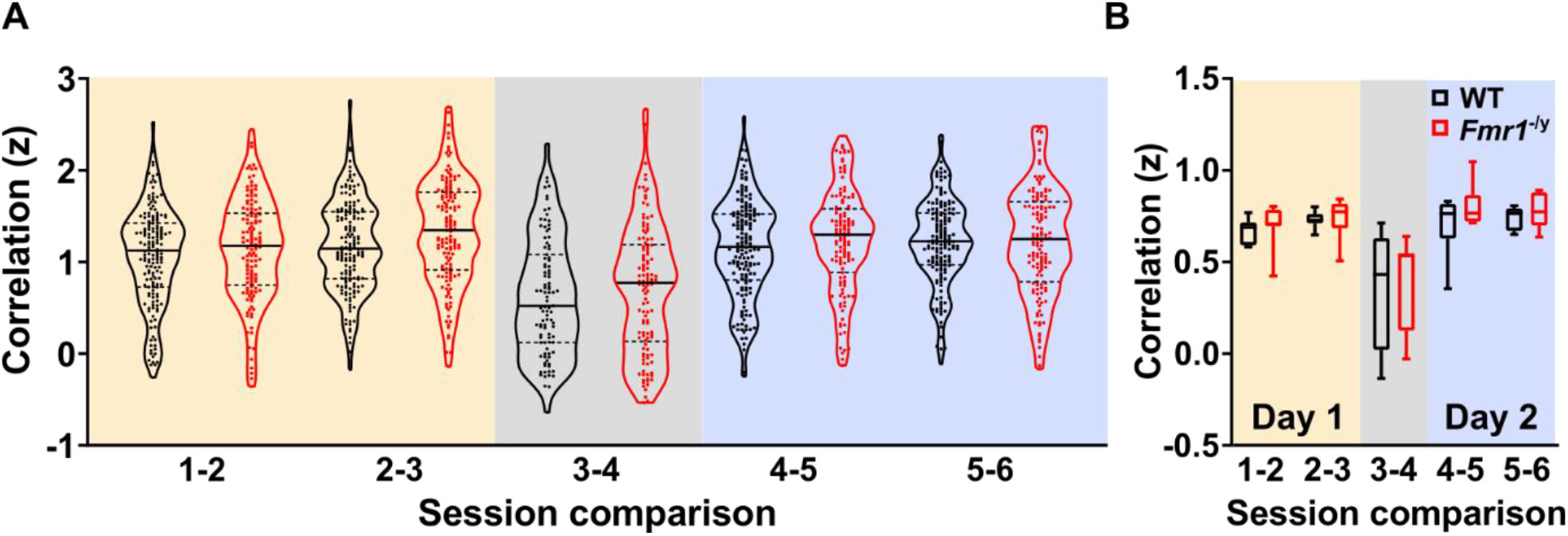
**(A)** Violin plots depicting the fisher z-transformed firing rate map correlation distributions across all five session comparisons for WT and *Fmr1*^*-/y*^ place cells. **(B)** Firing rate map correlation plotted and analysed at the rat average level. Two-way RM ANOVA: Comparison x genotype: F_(4,12)_=0.143, p=0.965; Genotype: F_(1,12)_=2.828, p=0.118; Comparison: F_(4,12)_=16.925, p<0.001. For box and violin plots in (**B**), the middle line represents rat median, upper and lower end of the boxes, and upper and lower line in the violins represent 95^th^ and 5^th^ percentile, error bars in box plots represent maximum and minimum values. N_WT_ = 7, N_KO_ =7. Pale yellow and pale purple backgrounds denote data from Day 1 and Day 2 respectively. Pale yellow and pale purple backgrounds denote data from Day 1 and Day 2 respectively, while grey denotes comparison between the last session on Day 2 (Session 3) and the first session on Day 2 (Session 4).

**Figure 3 – Supplement Table 1.**
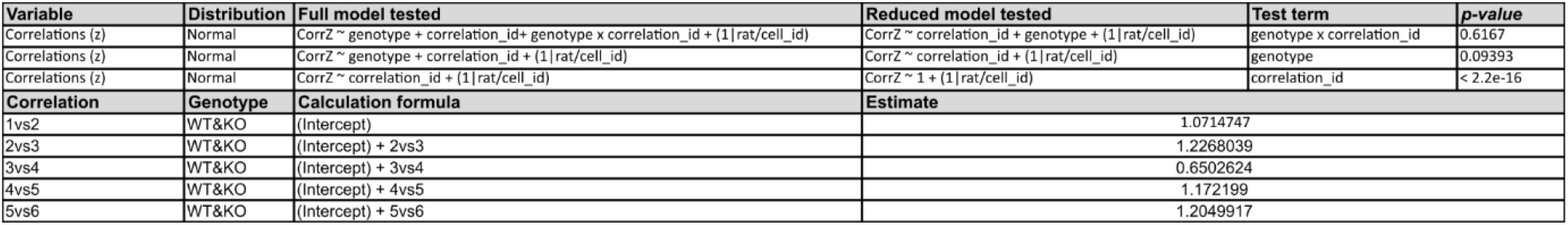
LME modelling estimates

**Figure 4 - Supplement 1.**
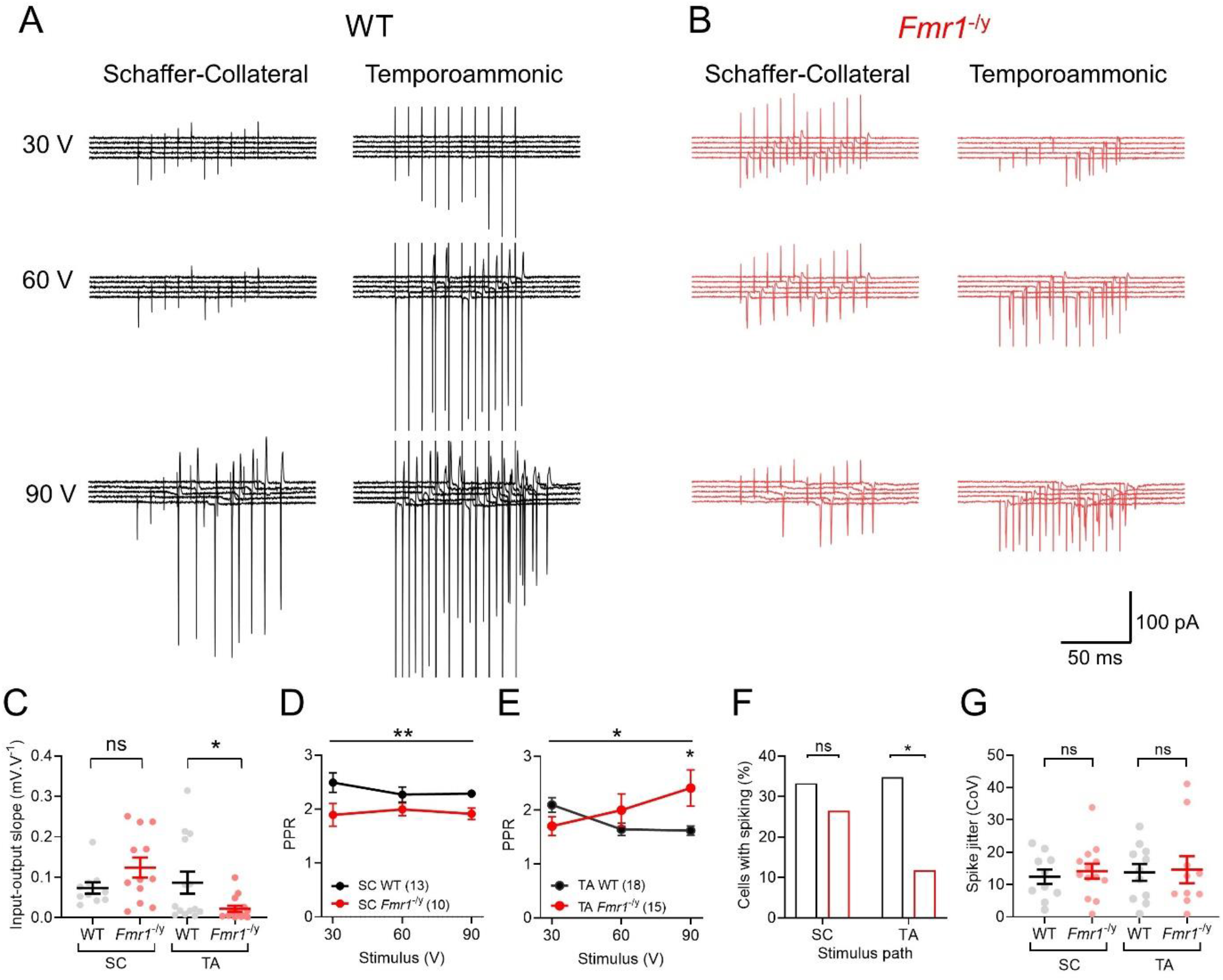
Genotype specific differences in synaptic and cellular recruitment at Schaffer-Collateral and Temporoammonic pathways. **A** Representative traces from a WT CA1 pyramidal neuron recorded in cell-attached configuration at the soma. 5 traces are shown at 30 V, 60 V, and 90 V stimulation delivered to the Schaffer-Collateral (left) or temporoammonic (right) paths in *str. radiatum* or *str. lacunosum-moleculare* respectively. **B** Representative traces performed under the same conditions as A, but for a pyramidal neuron from an *Fmr1*^*-/y*^ rat. **C** Quantification of the slope of input-output plots of EPSP amplitude at SC and TA paths in whole-cell recordings from CA1 pyramidal neurons. A greater slope is proportional to an increased input-output function. Data is shown as cell average from WT (black) and *Fmr1*^*-/y*^ (red) neurons, with individual neurons shown as open circles. Measurement of paired-pulse ratio (PPR) as derived from a 50 ms inter-pulse interval for all stimulation intensities from EPSP recordings are shown from SC (D) and TA (E) pathways. **F** Quantification of the number of neurons that responded with spiking to any stimulation intensity for WT and Fmr1 neurons, with respect to the stimulation pathway. **G** Measurement of the coefficient of variation (CoV) for the onset of action potentials driven by SC or TA afferents. All data is shown as mean ± SEM, except for **F**, where % of neurons is presented. Statistics shown: ns - p > 0.05 and * - p < 0.05, from 1-way ANOVA (**C**), GLMM (**D, E, G**) and Fisher’s exact test (**F**)

**Figure 4 - Supplement 2.**
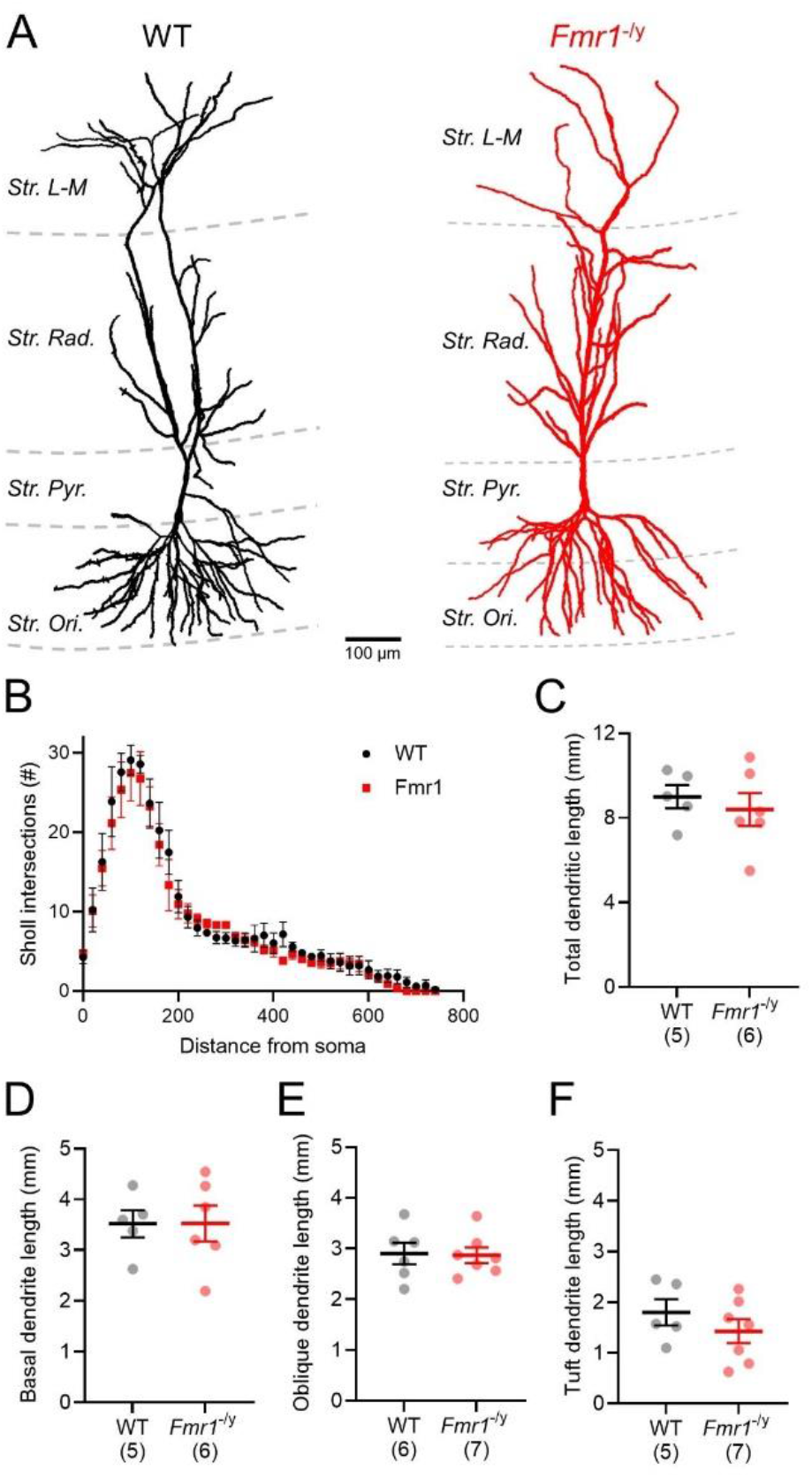
Typical morphology of CA1 pyramidal neurons from *Fmr1*^*-/y*^ rats. 3-dimensional reconstructions of representative CA1 pyramidal neurons from WT (black) and *Fmr1*-/y (red) rats, from neurons filled with biocytin during whole-cell patch-clamp recordings. Cells are shown with respect to hippocampal *stratum lacunosum-moleculare* (*Str. L-M*), *Radiatum* (*Str. Rad*.), *Pyramidale* (*Str. Pyr*.), and *Oriens* (*Str. Ori*), which are overlain as grey dashed lines. **B** Sholl analysis of reconstructed neurons, plotted as the average for all neurons recorded from WT (black, N=3 rats) and *Fmr1*^*-/y*^ (red, N=4) rats. **C** Measurement of total dendritic length of fully reconstructed neurons. Data of each rat are overlaid as filled circles. We observed no differences in the length of basal dendrites (**D**), apical oblique dendrites (**E**) or apical tuft dendrites (**F**), as measured between genotypes. Data from additional rats were included where basal dendrites were not filled sufficiently well for reconstruction. Data is shown as mean ± SEM with statistics from GLMM analysis.

**Figure 4 - Supplement 3.**
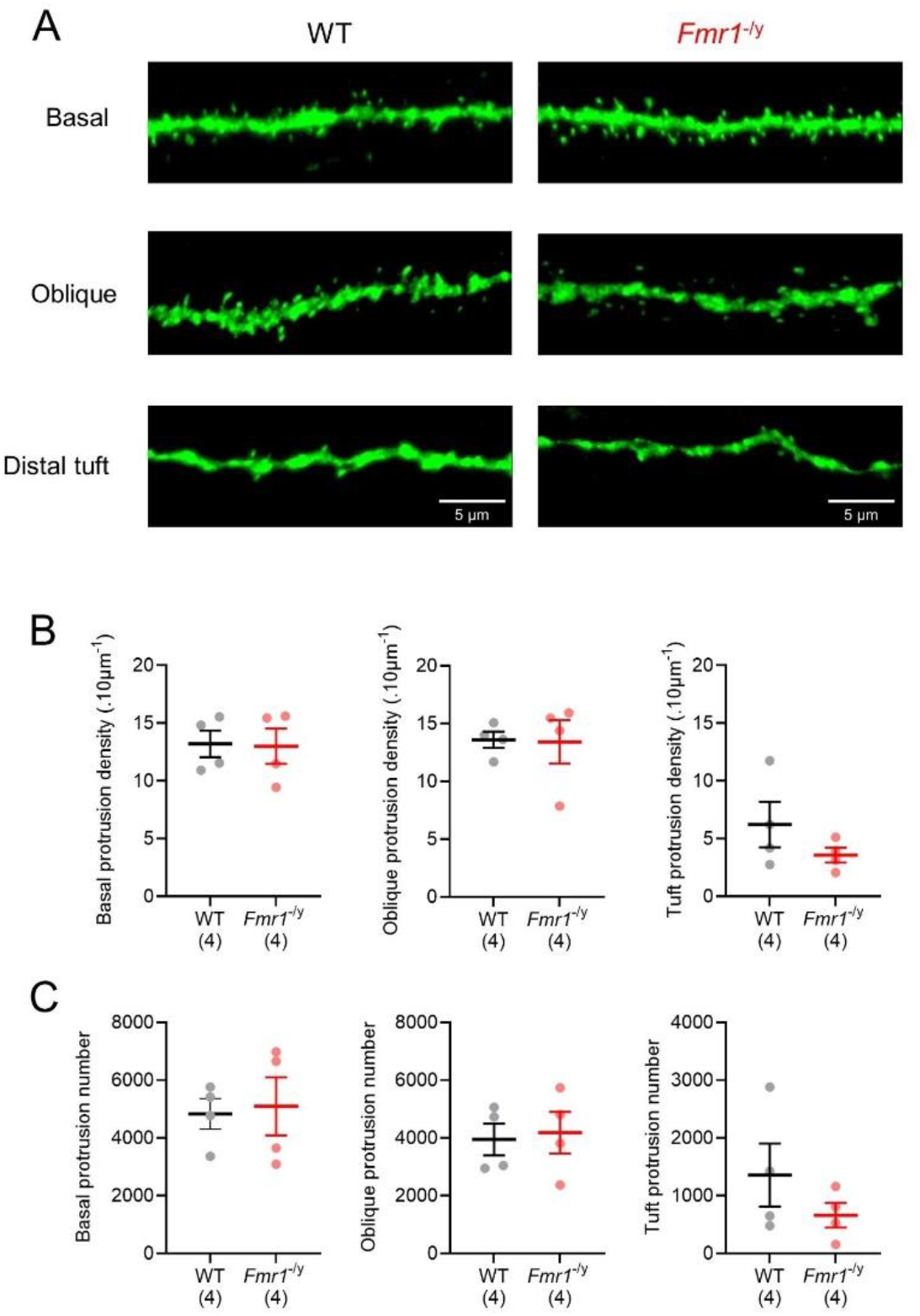
Reduced numbers of apical dendrite protrusions in CA1 pyramidal neurons from the *Fmr*^1-/y^ rat. Representative deconvolved and flattened confocal z-stacks from basal (upper), apical oblique (middle) and apical tuft (lower) dendrites of biocytin filled CA1 pyramidal neurons, from either wild-type (left) or *Fmr1*^-/y^ (right) rats. (**B**) Quantification of the density of protrusions from the same three dendritic compartments for wild-type (black, N=4) and *Fmr1*^-/y^ (N=4) rats. No difference between genotype was noted for any compartment (basal: *t*_(d.f. 6)_= 0.109, *p*=0.92; oblique: *t*_(d.f. 6)_= 0.089, *p*=0.93; apical tuft: *t*_(d.f. 6)_= 1.279, *p*=0.248; unpaired Student’s t-test). **(C)** Arithmetic sum of total protrusion number for the different compartments, based on the reconstructed neurons the dendrites were sampled from. No statistical difference was identified in any compartment, but a tendency towards reduced protrusion number was noted on the apical tuft dendrites (basal: *t*_(d.f. 6)_= 0.233, *p*=0.82; oblique: *t*_(d.f. 6)_= 0.258, *p*=0.81; apical tuft: *t*_(d.f. 6)_= 1.186, *p*=0.281; unpaired Student’s t-test). All means are superimposed with the results of individual rats.

**Figure 4 – Supplement Table 1.**
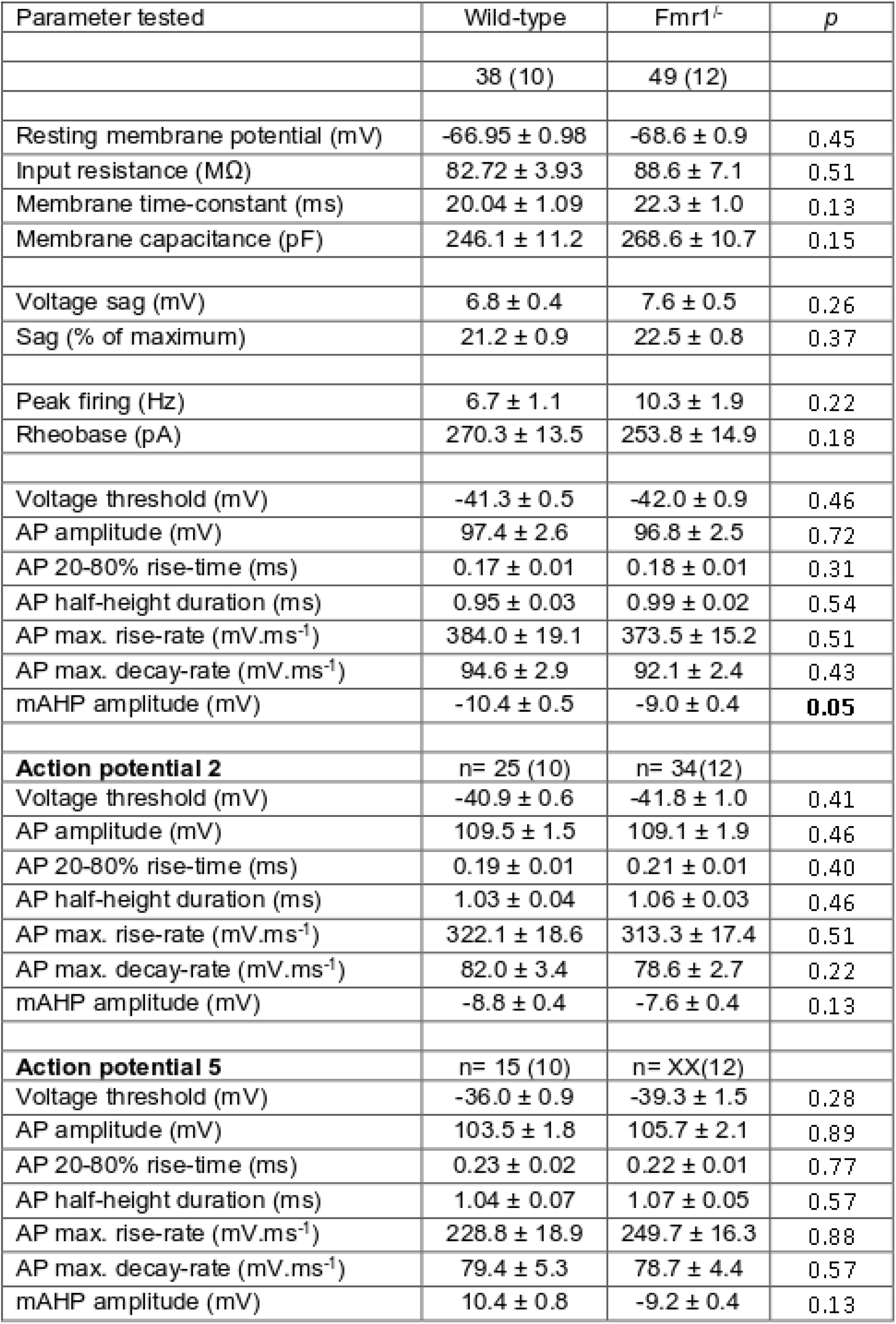
LME modelling estimates

**Figure 5 - Supplement 1.**
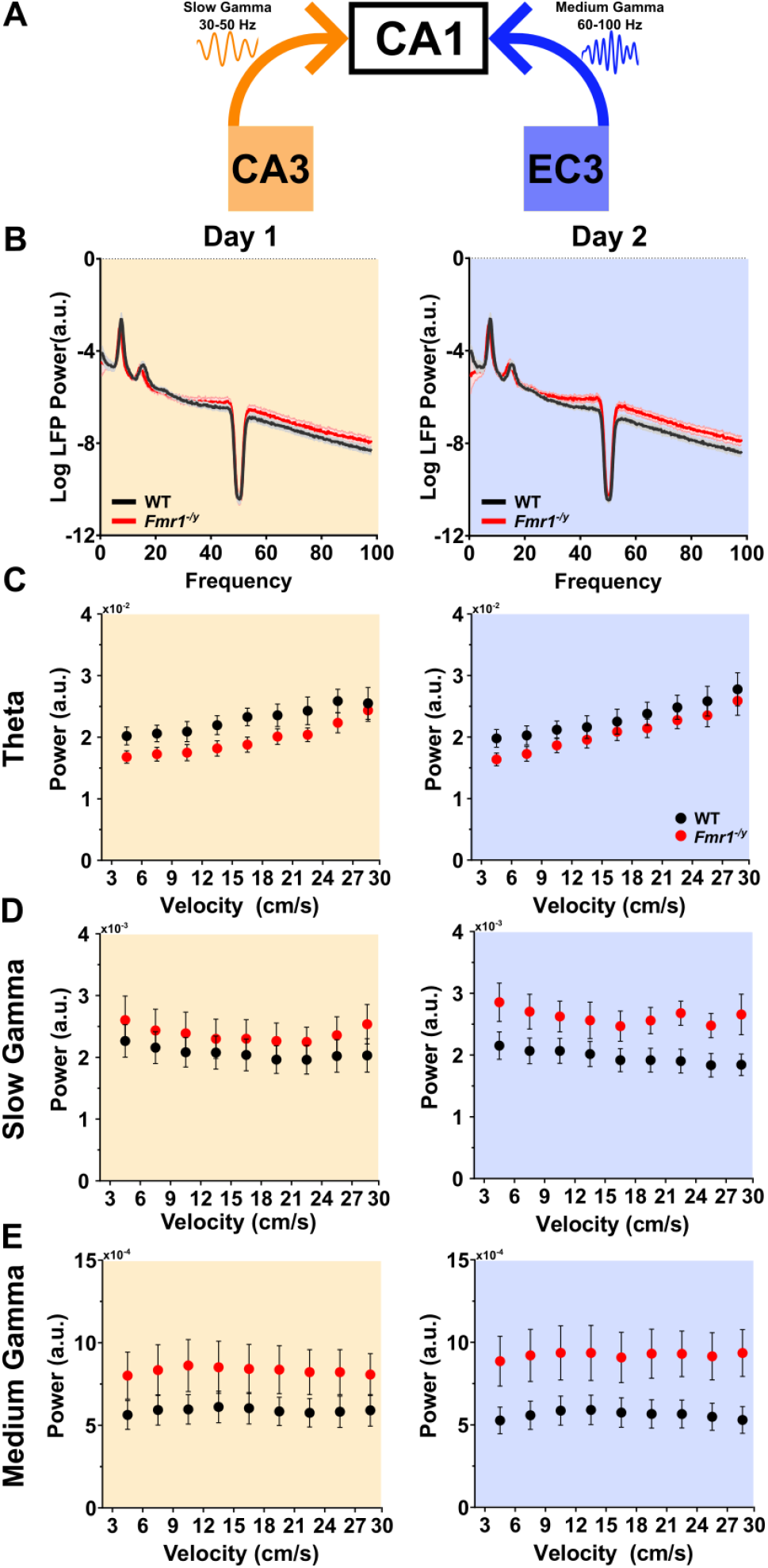
Power of hippocampal oscillatory activity and its modulation by animal velocity is not significantly different between WT and *Fmr1*^*-/y*^ rats. (**A**) Schematic depiction of the two major inputs to dorsal CA1. Inputs arriving from CA3 (yellow) are associated with slow gamma neural oscillations. Inputs from MEC3 (blue) are associated with medium gamma neural oscillations. (**B**) Mean LFP power spectra on Day 1 (left) and Day 2 (right) from WT and *Fmr1*^*-/y*^ rats during the first 4 s of each continuous recorded segment of movement following a period of immobility (>3 cm/s). Solid lines depict rat means; shaded areas depict SEM. Pale yellow and pale purple backgrounds denote data from Day 1 and Day 2 respectively. Theta. Three-way RM ANOVA: Velocity x genotype x day: F_(8,96)_=1.046 p=0.408; Genotype x day: F_(1,12)_=0.953, p=0.348; Velocity x genotype: F_(8,96)_=1.348, p=0.229; Genotype: F_(1,12)_=1.474, p=0.248; Day: F_(1,12)_=1.127, p=0.309, Velocity: F_(8,96)_=42.676, p<0.0001. **(C)** Slow gamma. Three-way RM ANOVA: Velocity x genotype x day: F_(8,96)_=0.577 p=0.794; Genotype x day: F_(1,12)_=3.605, p=0.082; Velocity x genotype: F_(8,96)_=1.517, p=0.161; Genotype: F_(1,12)_=1.350, p=0.267, Day: F_(1,12)_=1.247, p=0.286, Velocity: F_(8,96)_=3.127, p=0.003. **(D)** Medium gamma. Three-way RM ANOVA: Velocity x genotype x day: F_(8,96)_=0.919, p=0.524; Genotype x day: F_(1,12)_=6.813, p=0.023; Velocity x genotype: F_(8,96)_=0.833, p=0.576; Genotype: F_(1,12)_=2.702, p=0.126; Day: F_(1,12)_=1.291, p=0.156; Velocity: F_(8,96)_=2.472, p=0.018;). *Posthoc* One-way ANOVA Day1: F_(1,12)_=4.252, p=0.062; Day2: F_(1,12)_=1.240, p=0.287; WT: F_(1,6)_=0.406, p=0.548; *Fmr1*^*-/y*^: F_(1,6)_=16.416, p=0.007.

**Figure 5 - Supplement 2.**
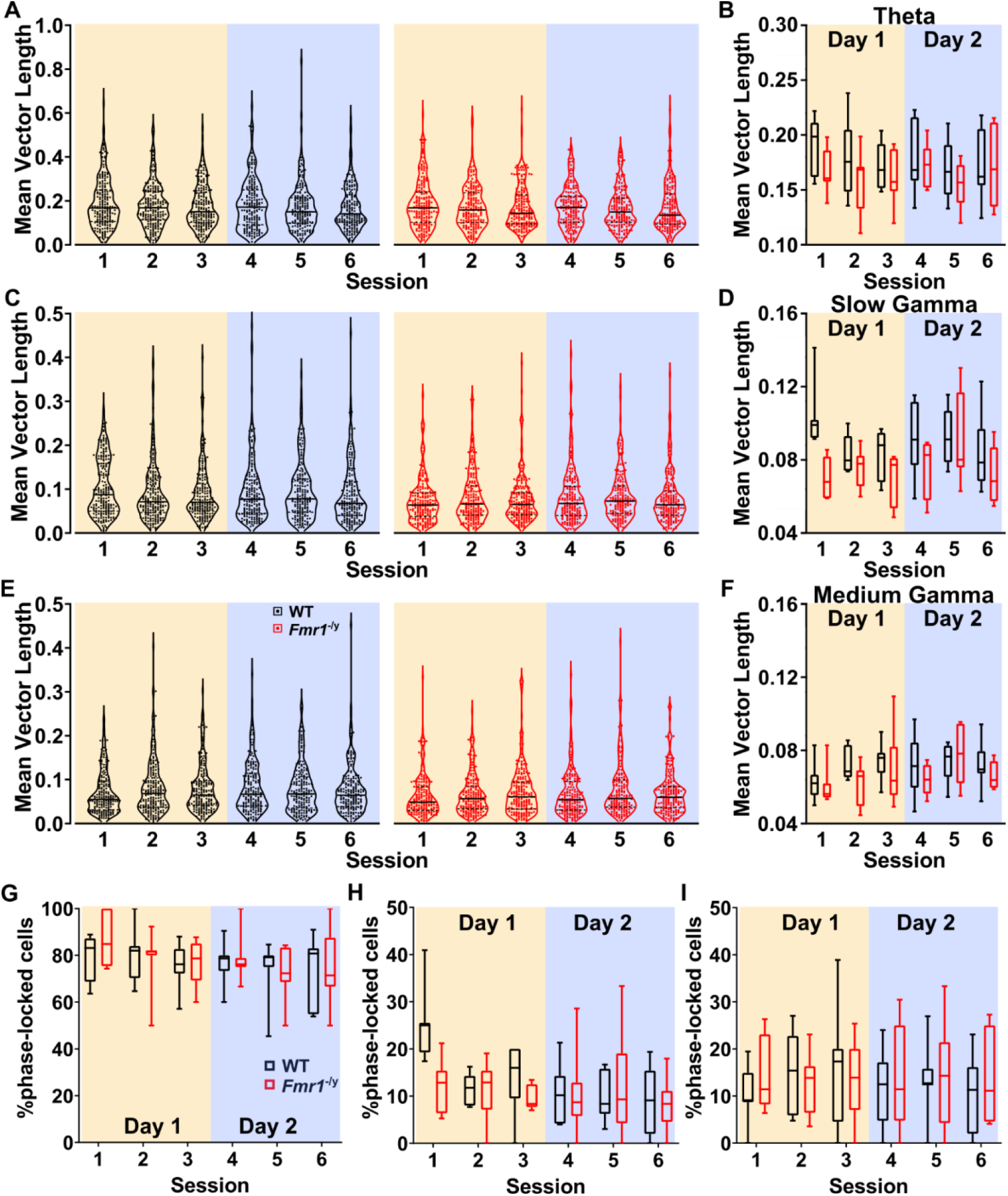
**(A)** Violin plots depicting the Theta MVL distributions across all six recording sessions for WT and *Fmr1*^*-/y*^ pyramidal neurons. **(B)** Theta MVL data plotted and analysed at the rat average level. Three-way RM ANOVA: Day x Session x genotype: F_(2,24)_=0.074, p=0.928; Day x Genotype: F_(1,12)_=0.585, p=0.459; Session x Genotype: F_(2,24)_=0.756, p=0.480; Genotype: F_(1,12)_=0.671, p=0.429; Day: F_(1,12)_=0.372, p=0.553; Session: F_(2,24)_=3.463, p=0.048. **(C-D)** Same as (**A**-**B**) for MVL Slow Gamma. Three-way RM ANOVA: Day x Session x genotype: F_(2,24)_=0.923, p=0.411; Day x Genotype: F_(1,12)_=1.492, p=0.245; Session x Genotype: F_(2,24)_=4.101, p=0.029; Genotype: F_(1,12)_=5.633, p=0.035; Day: F_(1,12)_=0.637, p=0.440; Session: F_(2,24)_=3.195, p=0.059. *Posthoc* tests Session1&4: WT vs *Fmr1*^*-/y*^ p=0.005; Session2&5: WT vs *Fmr1*^*-/y*^ p=0.553; Session3&6: WT vs *Fmr1*^*-/y*^ p=0.167. **(E-F)** Same as (**A**-**B**) for MVL Medium Gamma. Three-way RM ANOVA: Day x Session x genotype: F_(2,24)_=3.395, p=0.050; Day x Genotype: F_(1,12)_=0.618, p=0.447; Session x Genotype: F_(2,24)_=0.354, p=0.705; Genotype: F_(1,12)_=1.686, p=0.218; Day: F_(1,12)_=3.705, p=0.078; Session: F_(2,24)_=2.798, p=0.081. **(G)** Rat average values of percentages of Theta phase locked pyramidal neurons (Rayleigh p<0.05) across all six recording sessions. Three-way RM ANOVA: Day x Session x genotype: F_(2,24)_=0.025, p=0.976; Day x Genotype: F_(1,12)_=0.004, p=0.951; Session x Genotype: F_(2,24)_=0.014, p=0.986; Genotype: F_(1,12)_=1.830, p=0.201; Day: F_(1,12)_=1.035, p=0.329; Session: F_(2,24)_=0.353, p=0.706. **(H)** Same as (**G**) for Slow Gamma. Three-way RM ANOVA: Day x Session x genotype: F_(2,24)_=2.647, p=0.091; Day x Genotype: F_(1,12)_=2.433, p=0.145; Session x Genotype: F_(2,24)_=3.321, p=0.053; Genotype: F_(1,12)_=1.652, p=0.223; Day: F_(1,12)_=5.376, p=0.039; Session: F_(2,24)_=6.007, p=0.008. **(I)** Same as (**G**) for Medium Gamma. Three-way RM ANOVA: Day x Session x genotype: F_(2,24)_=0.492, p=0.617; Day x Genotype: F_(1,12)_=0.652, p=0.435; Session x Genotype: F_(2,24)_=0.068, p=0.934; Genotype: F_(1,12)_=0.009, p=0.928; Day: F_(1,12)_=0.001, p=0.990; Session: F_(2,24)_=0.660, p=0.526. For box and violin plots the middle line represents rat median, upper and lower end of the boxes, and upper and lower line in the violins represent 95^th^ and 5^th^ percentile, error bars in box plots represent maximum and minimum values. N_WT_ = 7, N_KO_ =7. Pale yellow and pale purple backgrounds denote data from Day 1 and Day 2 respectively.

**Figure 5 – Supplement Table 1.**
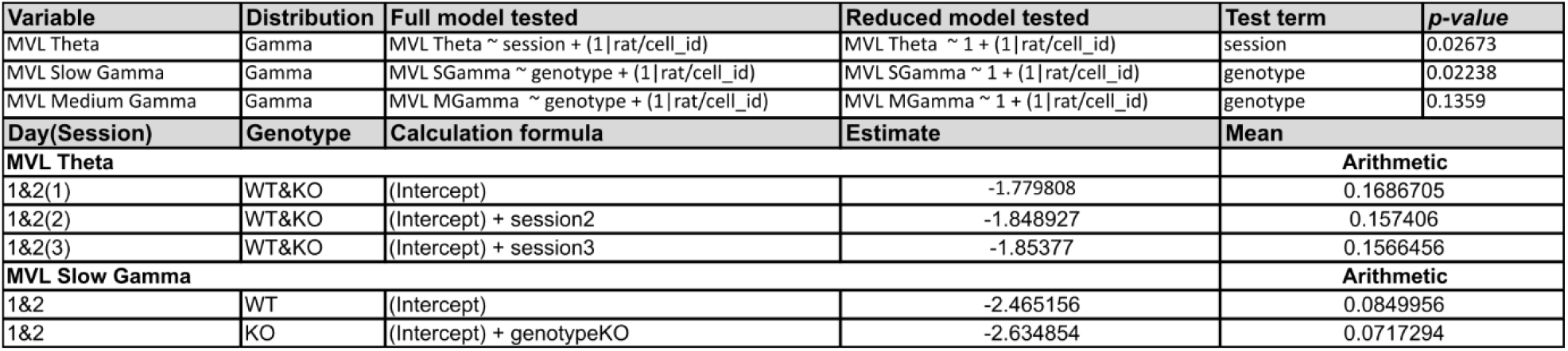
LME modelling estimates

